# Nerve injury disrupts temporal processing in the spinal cord dorsal horn through alterations in PV^+^ interneurons

**DOI:** 10.1101/2023.03.20.533541

**Authors:** Genelle Rankin, Anda M. Chirila, Alan J. Emanuel, Zihe Zhang, Clifford J. Woolf, Jan Drugowitsch, David D. Ginty

## Abstract

How mechanical allodynia following nerve injury is encoded in patterns of neural activity in the spinal cord dorsal horn (DH) is not known. We addressed this using the spared nerve injury model of neuropathic pain and *in vivo* electrophysiological recordings. Surprisingly, despite dramatic behavioral over-reactivity to mechanical stimuli following nerve injury, an overall increase in sensitivity or reactivity of DH neurons was not observed. We did, however, observe a marked decrease in correlated neural firing patterns, including the synchrony of mechanical stimulus-evoked firing, across the DH. Alterations in DH temporal firing patterns were recapitulated by silencing DH parvalbumin^+^ (PV^+^) inhibitory interneurons, previously implicated in mechanical allodynia, as were allodynic pain-like behaviors in mice. These findings reveal decorrelated DH network activity, driven by alterations in PV^+^ interneurons, as a prominent feature of neuropathic pain, and suggest that restoration of proper temporal activity is a potential treatment for chronic neuropathic pain.

## Introduction

An understanding of how the somatosensory system enables perception and reactivity to mechanical stimuli acting on the skin will guide the development of treatments for disorders of touch over-reactivity and mechanical pain. Chronic neuropathic pain, which can result from injury to the nervous system and is accompanied by painful reactivity to normally innocuous touch, called mechanical allodynia, afflicts between 3% and 17% of the global population^1–3^, and current therapies offer only moderate symptom amelioration and are associated with deleterious side effects^4, 5^. Defining the neurophysiological basis of mechanical allodynia has been a challenge, and previous findings related to the induction and expression of this form of painful touch have suggested sites of dysfunction in peripheral sensory neurons, the spinal cord, and the brain^1, 6–14^.

The first stage of integration of tactile signals flowing from the periphery is the spinal cord dorsal horn (DH). In the DH, axons of primary sensory neurons that innervate the skin and convey discrete streams of sensory information, including those in response to innocuous and noxious touch, synapse onto functionally diverse populations of interneurons as well as small populations of projection neurons^15–18^. The mechanosensory DH contains 10 or more interneuron subtypes, based on morphological, intrinsic physiological, molecular, and synaptic properties, at least four of which are inhibitory interneurons that collectively constitute ∼30% of the DH neuronal population^19–21^. The DH inhibitory interneurons mediate two principal forms of spinal inhibition: feedback inhibition via axo-axonic synapses onto primary afferent terminals, also known as presynaptic inhibition (PSI), and feed-forward inhibition (FFI) through axo-dendritic and axo-somatic synapses onto other interneurons and projection neurons^19, 21–23^. Spinal cord PSI and FFI are both necessary for normal output from the DH via projection neurons, including those projecting to higher order brain regions that underlie tactile perception and associated behavioral responses^15, 24–26^.

Changes in spontaneous and evoked activity of DH interneurons, particularly alterations in inhibitory interneurons leading to disinhibition, are a potential underlying cause of mechanical allodynia in neuropathic pain states^21, 27–30^, and indeed several DH interneuron subtypes have been implicated from morphological, behavioral, and *in vitro* physiological analyses^31–36^. However, the functional consequences of altered DH interneurons for population circuit dynamics of the intact spinal cord are not known. Thus, while synaptic inhibition can shape sensitivity, sensory tuning, spike timing, and network dynamics in other regions of the nervous system, including the cortex^37–42^, how alterations of synaptic inhibition in the DH in neuropathic pain states influence DH network activity, and the mechanisms and functional consequences, are unknown. Here, using *in vivo* multi-electrode array (MEA) recordings, we report that DH interneurons exhibit a temporal disorganization of spike patterns, but not hypersensitivity, in an injury-induced peripheral neuropathic pain state. Our findings also show that nerve injury and mechanical allodynia are associated with reduced PV^+^ interneuron activity, and that inhibition of these interneurons in uninjured mice both recapitulates decorrelated and desynchronized activity across the DH and causes concomitant pain-like behavior. Interestingly, alterations of other spinal cord inhibitory motifs, including disruption of GABA_A_R (GABA_A_ receptor)-dependent PSI, which has been theorized to underlie mechanical allodynia^43–45^, results in increased sensitivity, hyper-correlated DH activity, and behavioral over-reactivity, but not pain. Thus, a decrease in temporally correlated network activity in the DH, and not network over-reactivity, is a feature of the allodynic spinal cord, and restoration of proper network dynamics may be required to alleviate mechanical allodynia in chronic neuropathic pain states.

## Results

### *In vivo* spinal cord interneuron tuning in a mouse model of neuropathic pain

To assess *in vivo* physiological alterations in DH circuitry following spared nerve injury^46^ (SNI; Figure 1A) and the development of mechanical allodynia (Figure 1B), we characterized the response properties of individual DH interneurons by recording activity from dozens of neurons simultaneously using *in vivo* MEAs^15^. In urethane anesthetized mice, the region of lumbar spinal cord where neuronal receptive fields of lateral hindpaw glabrous skin are concentrated was targeted for recordings (Figure 1C) because this particular skin region remains innervated by sural nerve peripheral neurons after SNI and exhibits behavioral over-reactivity to normally innocuous mechanical stimuli. Force-controlled steps of indentation ranging from 1 to 75mN as well as gentle brush strokes were delivered to the lateral region of hindpaw glabrous skin. All recordings were done within 7 to 16 days post SNI surgery unless otherwise stated.

**Figure 1.**
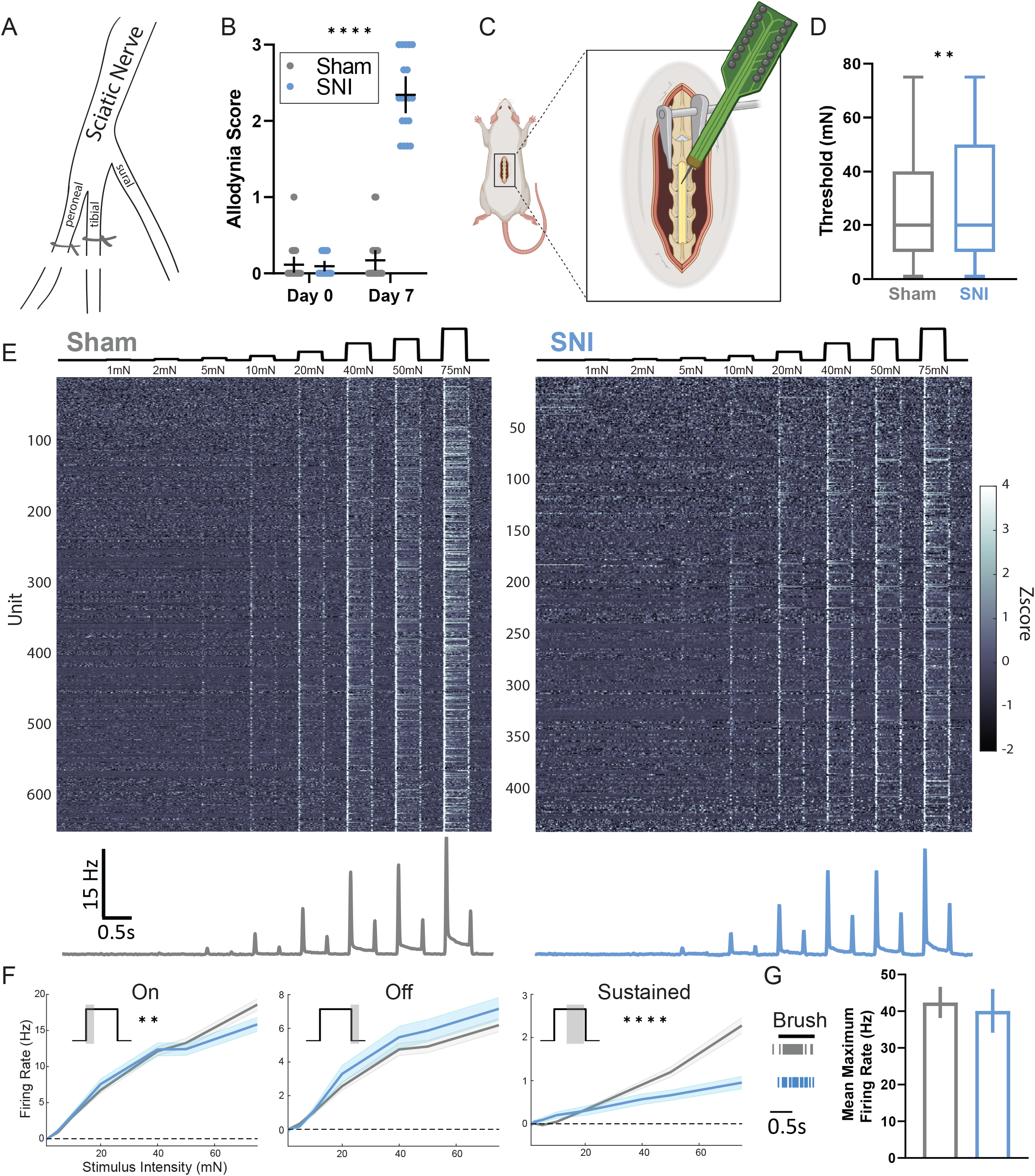
Mechanical allodynia following SNI is not associated with general physiologic over-reactivity across dorsal horn neurons. **A.** Diagram of the spared nerve injury model used to induce mechanical allodynia. The peroneal and tibial branches of the sciatic nerve are ligated and transected, sparing the sural branch, which innervated the lateral hindpaw. **B.** Dynamic allodynia score compared at Day 0 (prior to surgery) and Day 7 (post-surgery) between sham (N=22) and SNI (N=19) mice. Kruskal-Wallis H test with post-hoc Dunn’s test (H[3,82] = 51.37; p < 0.0001). SNI day 7 is significantly different from all other times points and conditions (****). **C.** Diagram of *in vivo* spinal cord MEA experimental setup. **D.** Distribution of indentation thresholds across DH neurons in sham (n=642, gray) and SNI (n=479, blue) mice. Mann-Whitney *U* test. **E.** Indentation responses for DH units in sham and SNI conditions. Top: force traces aligned to the heatmaps of Z-scored firing rates for each condition. Bottom: mean baseline-subtracted firing rate PSTHs. **F.** Average baseline-subtracted firing rates (±SEM) for DH units in sham and SNI groups at step indentation onset (On: 0-50ms after step onset), offset (Off: 0-50ms after step offset), and sustained (Sustained: 0-200ms before step offset) periods. On: two-way ANOVA [F (1,8952) = 9.024, p = 0.0027]. Sustained: two-way ANOVA [F (1,8952) = 36.72, p < 0.0001]. **G.** DH neurons responding to gentle brush strokes of the lateral hindpaw. Left: raster plot of an example sham (gray) and SNI (blue) neuron responding to brush. Right: average maximum brush evoked firing rates. Bars: mean. Error bars: 95% CI. Number of animals/ cells (N/n). **p < 0.01, ****p < 0.0001. See Table S1 for statistic details.

Using the dynamic brush assay^34^, robust nocifensive behaviors, including paw withdrawal, lateral kicking, and paw licking, were observed in all mice subjected to SNI but not sham surgery (Figures 1B and S1A). Sham and SNI mice were then used to assess the sensitivity and response properties of DH interneurons to test the hypothesis that SNI would cause a general increase in DH interneuron reactivity to the mechanical stimuli that cause pain. For this electrophysiological analysis, 642 randomly recorded single units (Figure S1B and S1C) spanning lamina I through V from 22 sham mice were obtained and compared to 479 units obtained from 19 SNI mice. Interestingly, although SNI animals exhibited dramatically increased behavioral responsivity to light touch stimuli (Figure 1B), no overall physiological over-reactivity was observed across the population of DH interneurons (Figure 1D-1G). In fact, contrary to our expectation, on average, DH interneurons exhibited a slight increase in their response thresholds to steps of static indentation following SNI (Figure 1D). We also generated tuning curves to investigate possible changes in firing rates to various components of the indentation steps. This analysis revealed that, as a population, DH interneurons in SNI mice did not exhibit changes in evoked firing rates during the OFF component of step indentations, but they did show decreased firing rates during the sustained portion of indentation steps and a small reduction in firing during the ON component (Figure 1E and 1F). In addition, the same brush stimulus used in the dynamic brush behavioral assay was used to stimulate the identical lateral hindpaw region while DH interneuron activity was recorded. There was no change in the maximum evoked firing rate to the brush stimulus between sham and SNI mice (Figure 1G). Thus, although dramatic behavioral over-reactivity to gentle touch was observed after nerve injury, as a population, DH interneurons did not exhibit physiological over-reactivity to tactile stimuli.

### Temporal activity patterns are disorganized after nerve injury

The lack of overall increased sensitivity and evoked activity in the DH of SNI mice led us to ask whether other aspects of DH circuit properties and firing patterns may be altered. The nature of our MEA recording configuration allowed for the activity of many neurons to be recorded simultaneously, and therefore whether and how temporal activity patterns change across the DH as a population in SNI mice was determined. To assess spike timing precision across populations of DH neurons, a population coupling metric was used to determine the extent to which an individual DH neuron’s spiking is correlated with other neurons in the population^47, 48^. Thus, by analyzing spike patterns across all simultaneously recorded units with 1ms time bins, the extent of synchronous population activity was determined during both indentation and brushing of the skin. A decrease in the synchrony of evoked spiking (see methods) during both indentation steps (Figure 2A) and brushing (Figure S2A) was observed in SNI mice compared to sham controls.

**Figure 2.**
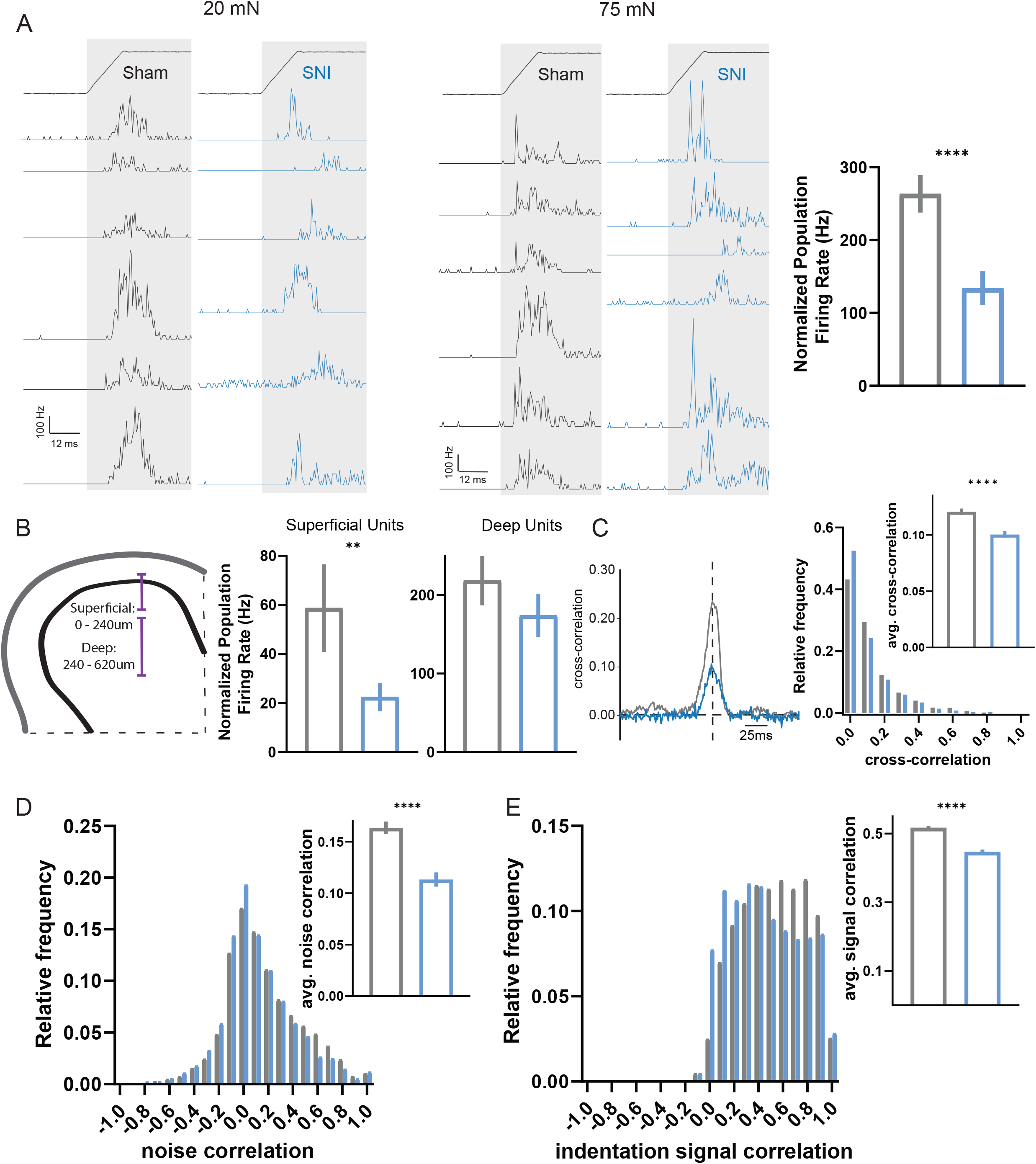
Temporal correlations of neuronal activity are altered after SNI. **A.** PSTHs (0.5ms bins) of simultaneously recorded units showing temporal alignment at indentation onset at 20mN (left) and 75mN (middle) in sham and SNI, followed by population coupling quantified as normalized population firing rate (right). **B.** Left: schematic of the DH subdivided into superficial and deep segments. Right: population coupling of superficial and deep units across conditions. **C.** Example cross-correlogram of one sham (gray) and SNI (blue) interneuron pair (left) followed by distribution of synchrony cross-correlations for pairs of DH neurons (at time lag = 0). Inset: average paired synchrony cross-correlations. **D.** Distribution of noise correlations for pairs of DH neurons. Inset: average noise correlations. **E.** Distribution of signal correlations for pairs of DH neurons. Inset: average signal correlations. Bars: mean. Error bars: 95% CI. Mann-Whitney *U* tests, *p < 0.05, **p < 0.01, ****p < 0.0001. See Table S1 for statistic details.

Because the superficial DH receives and processes information about high-threshold mechanical stimuli and noxious stimuli, as well as thermal and other stimuli, whereas the deep DH receives many inputs from primary sensory neurons that encode innocuous, light touch signals^49–51^, we next asked whether the altered synchrony of evoked spiking in SNI animals is observed across both superficial and deeper regions, or is more restricted. To test this, the DH was divided into superficial (units with recorded depths between 0 and ∼240 µm in the spinal cord) and deep (units with depths between ∼240 and 620 µm) regions, and synchronous population activity was calculated for units within these boundaries. After SNI, synchronous population activity within deep DH units was comparable to sham controls, however synchronous activity in the superficial DH was reduced by more than half (Figure 2B). To further assess the extent to which firing synchrony was altered after SNI, a range of bin sizes was used for the population coupling analysis. A deficit in synchronous firing was still observed when the bin size was expanded from 1ms to 3ms, but not 10ms (Figure S2B), thus constraining the timescale of desynchrony in the SNI animals to precise millisecond spike timing. Similar deficits were observed when calculating firing synchrony between pairs of neurons using spike cross-correlograms^52^ and computing the correlation at time lag 0 (Figure 2C). Thus, the precise synchrony of touch-evoked firing in neurons of the superficial DH is compromised in a neuropathic pain state.

As a complement to measuring spike-timing correlations across millisecond timescales, longer timescale activity correlations (across seconds) were examined in sham control and SNI animals to better understand network tuning and connectivity. Through spike count correlations during windows of spontaneous activity^52^, trial to trial variability between pairs of neurons (i.e., noise correlations) was assessed. SNI mice showed reduced noise correlations compared to sham controls (Figure 2D). These mice also displayed smaller pairwise signal correlations^52^, a measurement of tuning similarity, to both indentation and stroke of the hindpaw (Figures 2E and S2C). Together, the decreases in noise and signal correlations suggest that pairs of DH interneurons share fewer common inputs and are less similarly tuned to indentation steps and brush strokes, potentially indicative of a greater diversity in overall neural responses. Also of note, we observed a small but significant increase in the latency to first spike for the ON component of step indentations at higher forces as well as an increase in the jitter of the ON response (Figure S2D) in SNI mice. Moreover, disorganized population coupling across DH units was not observed immediately after nerve injury (4 hours post-surgery), which is prior to the development of mechanical allodynia (Figure S2I and S2J). Instead, deficits in population coupling and noise and signal correlations were observed in both early (7-16 days post SNI surgery) and later stages (28 to 35 days post SNI surgery) of the neuropathic pain state (Figures 2 and S2E-S2H), suggesting that disorganization of temporal firing patterns in the DH emerges at the same time as the behavioral pain response and persists throughout the transition from acute to chronic pain. Thus, a temporal disorganization of spiking in the DH, but not overall physiological hypersensitivity, occurs coincidently with the development of behavioral over-reactivity to tactile stimuli.

### PV^+^ interneuron activity is altered in mechanical allodynia

Previous studies of cortical circuitry have implicated fast-spiking inhibitory interneurons in the control of spike timing, neuronal tuning, and firing correlations^37^. Additionally, a deficit of DH inhibition has been proposed to underlie the development of neuropathic pain^27–36, 43^. These prior findings led us to hypothesize that dysfunction of DH fast-spiking inhibitory interneurons may lead to the alterations in temporal processing observed in SNI mice. To investigate possible abnormalities in DH inhibition following SNI, we first analyzed extracellular waveforms to dissect fast-spiking and regular spiking waveforms, as is routinely done in cortical datasets, to identify putative inhibitory neurons^53^. As opposed to the cortex, where both fast-spiking and regular spiking waveforms are readily observed (Figure S1D)^54, 55^, DH interneuron waveforms^15^ are more uniform and cannot be easily subdivided (Figure S1E). Using this extracellular waveform analysis, we observed no gross changes in DH waveforms between sham and SNI conditions (Figure S1F-S1I). We therefore turned to opto-genetic approaches to identify genetically distinct interneuron populations. All inhibitory interneurons were targeted using *Vgat^iCre^; R26^LSL-ChR^*^2^*^-YFP^* mice for identification using optical stimuli, and their responses in the spinal cord to mechanical stimuli were recorded *in vivo* using MEAs (Figure S3A-S3C). As a population, Vgat^+^ interneurons in SNI mice did not exhibit changes in response thresholds or evoked and spontaneous firing rates compared to sham controls (Figure S3D-S3G).

Since *Vgat^iCre^* labels all inhibitory interneurons, it remained possible that a small subset of inhibitory neurons may be altered after SNI, and these changes could be masked when analyzing the broader inhibitory interneuron population. Because of the net population reduction in sustained responses after SNI, we suspected that a fast-spiking inhibitory interneuron subtype known to have strong sustained responses to mechanical stimuli, the PV^+^ inhibitory interneurons of the DH^15^, may be affected in SNI mice. This population is of interest because *in vitro* and behavioral studies have suggested that PV^+^ interneuron intrinsic excitability is altered after SNI and that PV^+^ interneuron output and connectivity are necessary for behavioral over-reactivity to mechanical stimuli^31, 36, 56^. Although these prior findings implicated PV^+^ interneurons in neuropathic pain, whether evoked activity and sensory tuning of PV^+^ neurons change *in vivo* in neuropathic pain states remains unknown. Therefore, we opto-tagged PV^+^ interneurons using *PV^Cre^; R26^LSL-ChR^*^2^*^-YFP^* mice for *in vivo* electrophysiological recordings to assess their physiological responses to tactile stimuli following SNI (Figure 3A and 3B). It is noteworthy that *PV^Cre^* labels subsets of excitatory and inhibitory neurons; approximately 70-95% of DH PV^+^ cells are co-labelled with inhibitory markers^19, 36^ (Figure 3A). After SNI, PV^+^ interneurons did not exhibit changes in their response thresholds or spontaneous firing rates (Figure 3C and 3D). However, these interneurons displayed decreased firing to the sustained and OFF portions of step indentations, as well as reduced firing to brushing the skin (Figure 3E-3G). PV^+^ interneurons also showed decreased noise correlations with other simultaneously recorded units suggesting a reduction in PV^+^ interneuron connectivity after SNI (Figure 3H), consistent with a previous anatomical study^36^, while no changes in signal correlations were observed (Figure 3H). In a complementary set of experiments, CCK^+^ interneurons, a broad population of excitatory neurons implicated in neuropathic pain via their connections with corticospinal tract axons^57^, were also examined after SNI using *CCK^iCre^; R26^LSL-ChR2-YFP^* mice. These excitatory CCK^+^ interneurons showed increased responsivity at the OFF portion of the step indentations (Figure S3H-S3K) suggesting CCK^+^ interneurons may contribute to circuit disruption following nerve injury. This increased firing is consistent with a decrease in PV-mediated inhibition, since PV^+^ interneurons, which show reduced evoked firing following nerve injury (Figure 3F), form synapses onto at least some CCK^+^ interneurons ^36^.

**Figure 3.**
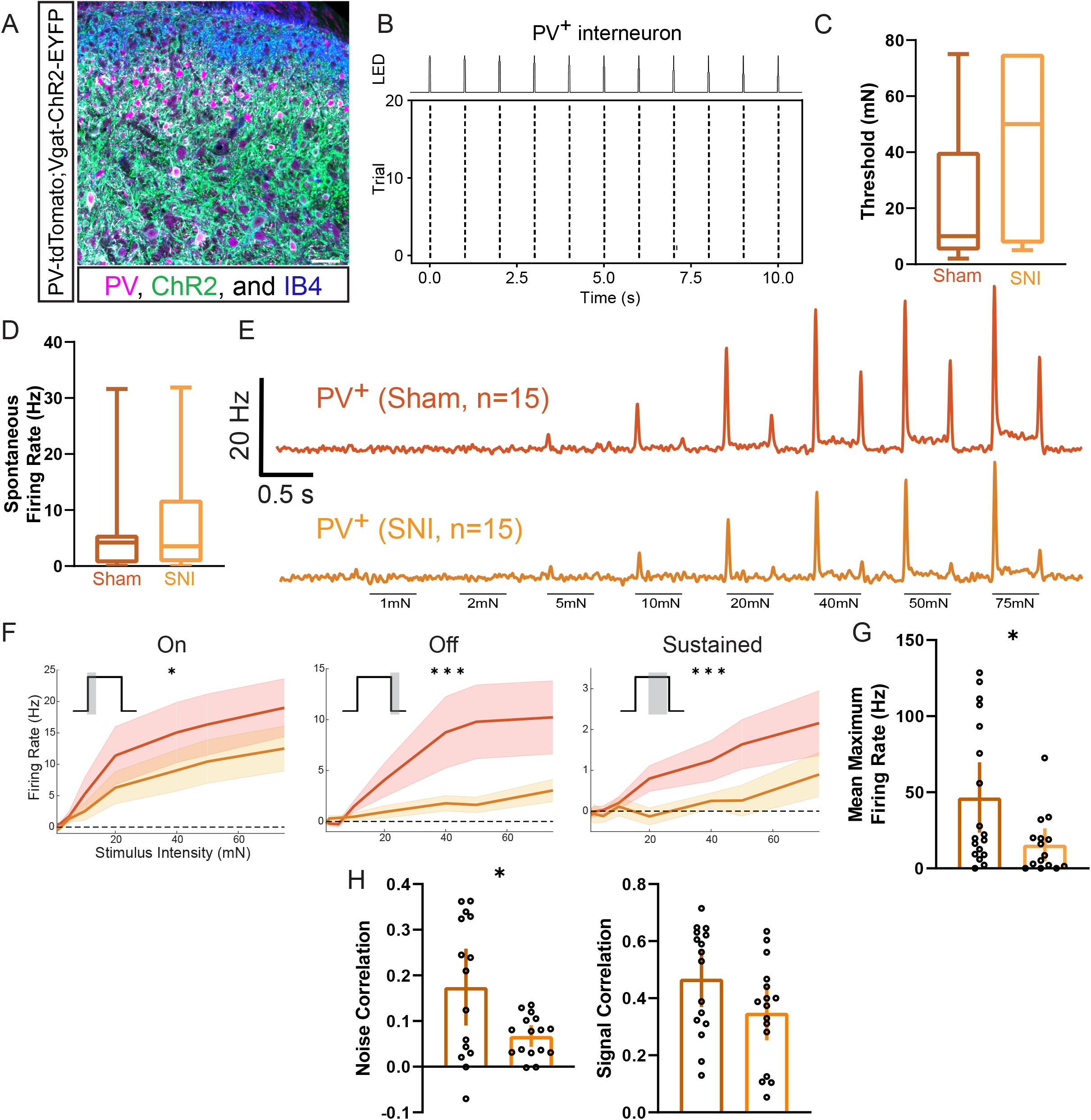
Dorsal horn PV^+^ interneuron activity is decreased in mice following SNI. **A.** Coronal spinal cord section showing PV^+^ interneurons in comparison to all Vgat^+^ cells (PV-tdTomato;Vgat-ChR2-EYFP mouse). PV^+^ neurons; magenta, Vgat-ChR2^+^ neurons; green, IB_4_ binding; blue (labels lamina IIi). Scale bar: 50 µm. **B.** Raster plot of light-evoked spikes in a PV^+^ interneuron. **C.** Indentation thresholds across PV^+^ neurons in sham (N=3, n=15) and SNI (N=3, n=15) mice. **D.** Spontaneous firing rates of PV^+^ interneurons in sham (N=3, n=15) and SNI (N=3, n=15) mice. **E.** Mean baseline-subtracted firing rate PSTHs for sham and SNI PV^+^ neurons. **F.** Average baseline-subtracted firing rates (±SEM) for PV^+^ neurons in sham and SNI groups at step indentation On, Off, and Sustained periods. On: two-way ANOVA [F (1,240) = 5.602 p = 0.0187]. Off: two-way ANOVA [F (1,240) = 14.59, p = 0.0002]. Sustained: two-way ANOVA [F (1,240) = 11.41, p = 0.0009]. **G.** Sham and SNI PV^+^ interneurons average maximum brush evoked firing rates. Unpaired t-test. **H.** Noise (left) and indentation signal (right) correlations between PV^+^ interneurons and neighboring cells in sham and SNI mice. Unpaired t-test. Bars: mean. Error bars: 95% CI. Number of animals/ cells (N/n). *p < 0.05, **p < 0.01, ***p < 0.001. See Table S1 for statistic details.

### PV^+^ interneurons control temporal activity patterns across spinal cord neurons and mechanical allodynia following nerve injury

The marked reduction in PV^+^ interneuron firing following SNI raised the possibility that PV^+^ interneurons control DH temporal dynamics. To test this, PV^+^ interneurons were silenced with tetanus toxin using *PV^Cre^;Lbx1^FlpO^;RC::PFtox* mice, and spike timing precision with population coupling and paired firing synchrony was measured using *in vivo* MEA recordings. Remarkably, as observed in SNI mice, evoked population coupling to both indentation (Figure 4A) and brush (Figure S4A) was markedly decreased in PV-silenced mice. Moreover, as in SNI mice, these coupling deficits were most pronounced in the superficial DH (Figure 4B). Also similar to findings with SNI mice, the deficits in population coupling in PV-silenced mice diminished as bin sizes increased, thus resolving the synchrony deficits to millisecond spike timing (Figure S4B). Thresholds of DH interneurons remained unchanged in response to PV-silencing (Figure S4C), however, as in SNI, paired synchronous firing and noise and signal correlations were decreased in PV-silenced mice (Figures 4C-4D and S4D), suggesting that DH interneurons share fewer common inputs and that evoked responses to mechanical stimuli are more diverse when PV^+^ mediated synaptic transmission is compromised. Thus, PV^+^ interneurons exhibit reduced activity under SNI conditions, and silencing PV^+^ interneurons recapitulates the marked disorganization of temporal firing patterns in the superficial DH observed in the neuropathic pain state (Figure S4E-S4G).

**Figure 4.**
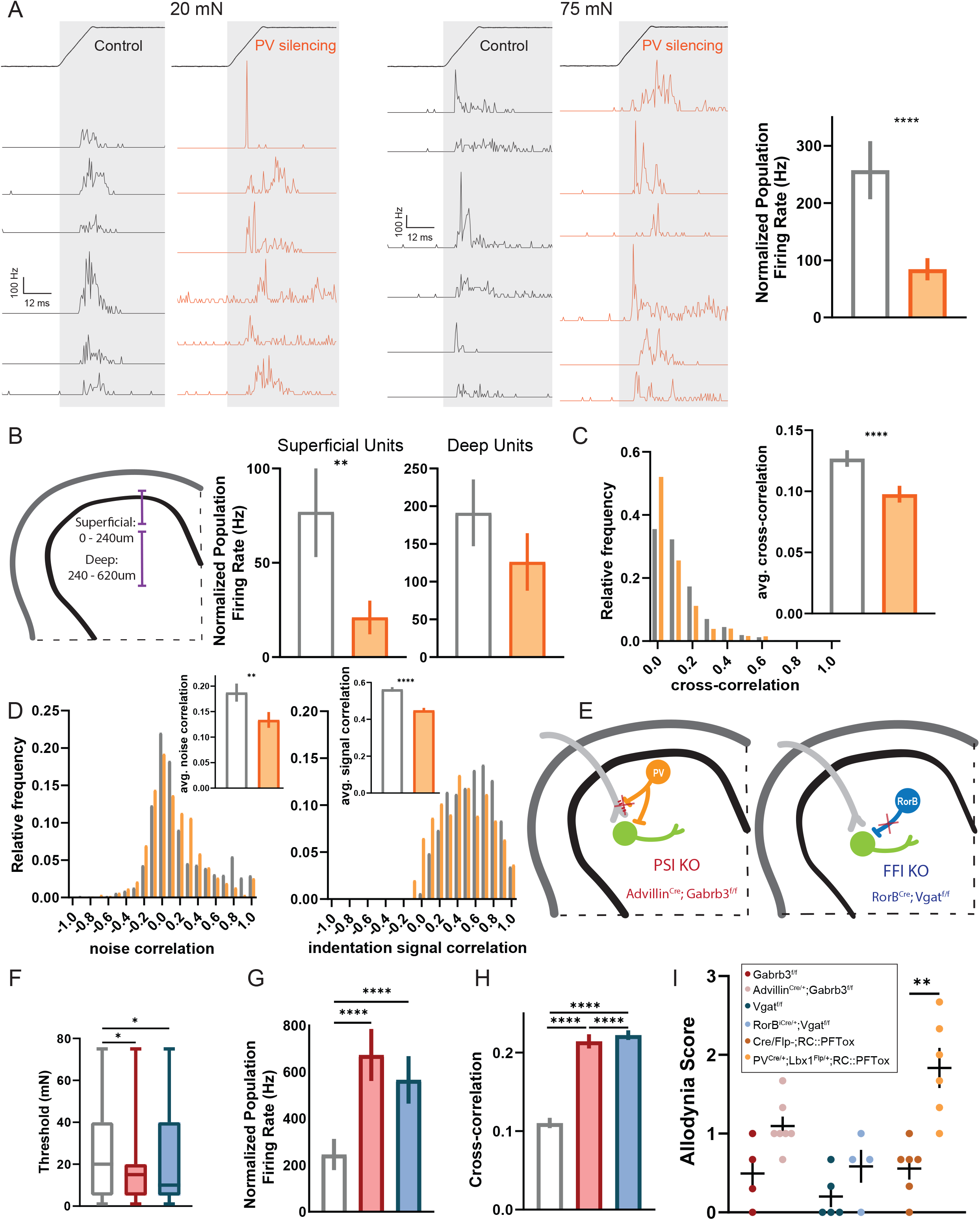
Deficits in temporally correlated activity across the DH and allodynia-like behavior after silencing PV^+^ interneurons. **A.** PSTHs (0.5ms bins) of simultaneously recorded units showing temporal alignment at indentation onset at 20mN (left) and 75mN (middle) in control (*PV^Cre^;RC::PFtox*, N=4) and PV-silencing (*PV^Cre^;Lbx1^FlpO^;RC::PFtox*, N=4) conditions, followed by population coupling quantified as normalized population firing rate (right). Mann-Whitney *U* test. **B.** Left: schematic of superficial and deep DH segments. Right: population coupling of superficial and deep units across conditions. Mann-Whitney *U* tests. **C.** Distribution of synchrony cross-correlations for pairs of DH neurons in controls (white bars) and mutants (orange bars). Inset: average paired cross-correlations. Mann-Whitney *U* test. **D.** Distributions of noise (left) and signal (right) correlations for pairs of DH neurons. Insets: average noise and signal correlations in controls (white bars) and mutants (orange bars). Mann-Whitney *U* tests. **E.** Diagram of genetic strategies to silence all presynaptic DH inhibition (*Avil^Cre^;Gabrb3^f/f^*; PSI KO) and Rorβ-mediated feedforward inhibition (*Ror*β*^iCre^;Vgat^f/f^*; FFI KO). **F.** Distribution of indentation thresholds between controls (*Vgat^f/f^* or *Gabrb3^f/f^*, white, N=3), PSI KOs (red, N=3), and FFI KOs (blue, N=3). One-way ANOVA with post-hoc Tukey’s test [F (2, 311) = 4.606; p = 0.0107]. **G.** Population coupling for each condition. Kruskal-Wallis H test with post-hoc Dunn’s test (H[2,314] = 42.30; p < 0.0001). **H.** Synchrony cross-correlations for neuron pairs. Kruskal-Wallis H test with post-hoc Dunn’s test (H[2, 6099] = 757.3; p < 0.0001). **I.** Dynamic allodynia score compared at baseline (no surgery). Mean and ±SEM plotted. Mann-Whitney *U* test. Bars: mean. Error bars: 95% CI. Number of animals (N). **p < 0.01, ***p < 0.001, ****p < 0.0001. See Table S1 for experimental details.

We next asked how PV^+^ interneurons contribute to synchrony in DH firing patterns. Inhibitory PV^+^ interneurons form two types of synapses in the DH: axo-dendritic synapses, contributing to feed-forward inhibition (FFI), and axo-axonic synapses, contributing to GABA_A_R-dependent pre-synaptic inhibition (PSI) of primary afferent terminals^19, 31, 58^ (Figure S4H). To determine whether one or both forms of PV-mediated inhibition is responsible for the coordinated activity deficits observed following PV^+^ interneuron-silencing, and potentially underlying the coordinated activity changes in SNI, DH neuron activity was recorded in mice lacking GABA_A_ receptors on primary afferents, which are necessary for GABA_A_R-dependent PSI^25, 26^ (*Avil^Cre^;Gabrb3^f/f^* mice, Figure 4E). Using the same set of coordinated activity metrics, we observed increased population coupling, paired synchronous firing, and signal correlations (Figures 4G-4H and S4I) as well as decreased indentation thresholds (Figure 4F) in *Avil^Cre^;Gabrb3^f/f^* mice, precisely the opposite of that observed following SNI or PV^+^ interneuron silencing. These findings indicate that ablating PSI alone does not recapitulate the deficit in temporal correlations observed following PV^+^ interneuron-silencing and SNI. Thus, either PV^+^ interneuron axo-dendritic connections must be compromised or both axo-axonic and axo-dendritic connections must be compromised to cause the alterations in temporal correlation in SNI and PV^+^ interneuron-silenced mice. Although we were unable to selectively disrupt the FFI function of PV^+^ interneurons, we used *Ror*β*^iCre^;Vgat^f/f^* mice in which Rorβ inhibitory interneurons are silenced, to ask whether other forms of FFI disruption cause similar temporal changes across the DH (Figure 4E). Rorβ interneurons make axo-dendritic synapses in the deep DH (laminae IIiv – IV)^19^ and provide the majority of mechanically evoked FFI of post-synaptic dorsal column projection neurons^15^. These experiments using *Ror*β*^iCre^;Vgat^f/f^* mice revealed changes in sensitivity and correlated evoked activity (Figures 4G-4H and S4I) similar to the PSI disruption model but distinct from the temporal dysregulation seen in SNI and PV^+^ interneuron-silencing. These findings thus point to a unique role for PV^+^ DH interneurons in coordinated network activity in the DH via their contribution to FFI, or a combination of FFI and PSI, but not PSI alone.

The requirement of PV^+^ signaling for normal temporal patterns of activity in the superficial DH, in conjunction with our findings that both PV^+^ interneuron activity and correlated activity are markedly reduced in SNI, led us to explore the extent to which silencing PV^+^ interneurons, disrupting GABA_A_R-dependent PSI, or disrupting Rorβ-mediated FFI, leads to a mechanical allodynia similar to SNI. Thus, using the dynamic brush assay to test for mechanical allodynia, PV-silenced, PSI-disrupted, and Rorβ-mediated FFI-disrupted mice were evaluated. PV^+^ interneuron-silenced mice exhibited an allodynia comparable to SNI (Figure 4I). This finding is consistent with prior results showing that ablating PV^+^ interneurons increased punctate mechanical sensitivity^36, 56^. On the other hand, disruption of either PSI or Rorβ-mediated FFI, both of which lead to a dramatic increased physiological reactivity to step indentations^15^ and increased correlated activity in the DH (Figures 4G-4H and S4I), caused behavioral over-reactivity to light touch but not pain-like behaviors reflected by lateral kicking and licking of the contacted paw (Figure 4I)^15, 24–26, 59^. Taken together, these findings suggest that decorrelated activity at the population level in the DH, resulting from altered PV^+^ interneuron activity, and not a generalized increased in evoked reactivity across the DH, underlies mechanical allodynia in a peripheral nerve injury-induced neuropathic pain state.

## Discussion

Using *in vivo* multi-unit electrophysiology, genetic labeling, and network-level activity analyses we aimed to identify physiological signatures that represent normal DH circuit function as well as circuit level dysfunction underlying mechanical allodynia associated with neuropathic pain. Our findings suggest that a deficit in coordinated activity, including temporal misalignment of touch-evoked DH interneuron spiking, and not general over-reactivity to tactile stimuli, is a characteristic feature of the allodynic spinal cord. Additionally, we found that PV^+^ interneurons control rapid (millisecond time scale) temporal processing in the DH, and that reduced activity of these interneurons in the allodynic spinal cord is responsible for aberrant temporal processing of tactile signals. We propose that, following nerve injury, a reduction in DH PV^+^ fast-spiking inhibitory interneuron activity underlies deficits in the synchrony of touch evoked spiking in the DH, which produces mechanical allodynia.

### Decorrelated network activity, not generalized over-reactivity, in the dorsal horn is observed in a neuropathic pain model

After SNI, animals exhibit nocifensive behaviors in response to normally innocuous stimuli, and yet our *in vivo* electrophysiological recordings showed a lack of increase in sensitivity or evoked firing across the general DH population. In fact, overall firing levels decreased during the sustained portion of step indentations in SNI animals. This suggests that the increased behavioral output is not directly linked to an overall increase in evoked spiking across the DH, as we had expected, and that other facets of neuronal processing are likely altered to account for the dramatic shift in behavioral reactivity. Indeed, we found that coordinated population activity and synchronous firing across DH neurons are markedly decreased after SNI. This change in synchronous population activity was not observed 4 hours post nerve injury but only arose over days, coincident with the development of pain-like behavioral responses to light touch.

After SNI, individual interneurons of the DH are less coupled to the total population activity, compared to DH interneurons of control mice, meaning that there are more neurons that are ‘soloists’, less influenced by population-wide events compared to a predominance of ‘chorister’ neurons observed in the control DH. Whether and how this loss of broad, synchronous population activity could lead to altered behavioral reactivity or perception of tactile stimuli is not known. Synchrony is often proposed to enable efficient information transfer from one brain region to another. One possibility is that desynchronized responses in the DH results in an incomplete representation of tactile features in downstream brain regions and therefore disables the ability of the CNS to decode or match with internal predictions of sensory experiences, as posited by the theory of predictive coding^60^. In fact, we observed a decrease in the similarity of neuronal tuning between pairs of neurons after SNI (decreased signal correlations). This suggests an increase in the variety of responses even to simple stimuli, and an expansion of the range of activity patterns emanating from the spinal cord to higher order regions; such novel physiological response patterns could lead to “misinterpretation” of light touch stimuli as being noxious.

It is also possible that the observed disorganization of spike timing in the neuropathic pain state allows for signals to propagate to DH projection neurons that would normally be blocked or shunted by precisely timed inhibitory connections. Such unchecked signals arising from primary afferents or neighboring DH interneurons could result in altered sensitivity, evoked firing, latency to respond, and signal-to-noise ratios, both at the level of individual neurons and as a population. For example, it is possible that synchronized inhibition is required to prevent low-threshold mechanoreceptor (LTMR) inputs from driving pain circuits and that this is lost following nerve injury, resulting in augmented DH projection neuron responses and nocifensive behavioral responses to tactile stimuli. Tests of these and other models will require probing changes in responses, including synchronous responses, across DH projection neuron populations after nerve injury; this will be challenging however, because superficial and deep DH projection neurons are relatively few in number and heterogenous in both tuning properties and genetic identity^15, 17–19, 61, 62^. Future goals will be to assess responses of DH output neurons with the population measurements used here and determine the degree to which PV^+^ inhibitory interneurons and the synchronization of DH firing patterns shape projection neuron responses.

### Dysfunction of distinct dorsal horn inhibitory motifs can drive tactile over-reactivity

To address the basis for loss of population coupling following SNI, we manipulated three types of DH inhibitory circuit motifs and found that these alterations yielded both distinct electrophysiological changes across the DH and different behavioral manifestations. First, we silenced DH PV^+^ interneurons blocking their contributions to both feed-forward and feedback inhibition. This manipulation led to disorganized, desynchronized firing, predominantly in the superficial DH, and increased allodynic behaviors, mimicking the population activity alterations and behavioral over-reactivity observed following SNI. Consistent with this, *in vivo* opto-tagging experiments showed that, following nerve injury, PV^+^ interneurons exhibit decreased evoked firing. Together, these findings suggest that alterations in PV^+^ interneurons underlie the uncoordinated population activity in the DH following nerve injury. It is interesting that alterations in PV^+^ interneuron activity in the cortex have also been linked to changes in neuronal activity correlations and spike timing^37–42^, indicating that PV^+^ inhibitory interneurons in at least two CNS regions coordinate the temporal dynamics of circuit function. We suspect that this curious parallel reflects the need for fast-spiking interneurons to coordinate synchrony across CNS regions, and that calcium binding proteins such as PV are expressed in fast-spiking neurons to buffer high levels of free, ionized calcium. Nevertheless, since SNI causes a reduction in touch-evoked excitation of DH PV^+^ interneurons *in vivo* and intrinsic excitability and homeostatic plasticity measured *in vitro*^31, 56^, future studies to investigate the basis of their altered physiological properties are needed.

Finally, it is noteworthy that while SNI and silencing PV^+^ interneurons caused reduced synchronous firing in the DH and concomitant mechanical allodynia, but little to no change in overall physiological sensitivity across the DH as a whole, silencing either GABA_A_R-dependent PSI or Rorβ-mediated FFI led to *increased* sensitivity and reactivity across the DH and *enhanced* synchronous firing, but not mechanical allodynia. Thus, opposing changes in spiking precision and correlated activity are associated with distinct behaviors, further implicating temporally decorrelated activity, and not general over-reactivity of DH interneurons, as the culprit driving mechanical allodynia following nerve injury. Our findings, which emphasize the importance of temporal encoding in touch signals in the DH, lead us to suggest that approaches that reinstate normal patterns of synchronous firing across DH interneurons may help restore normal behavioral reactivity to tactile stimuli and perception following nerve injury.

## Acknowledgements

We thank Aniqa Tasnim for assistance with histology, Ofer Mazor and Pavel Gorelik from the HMS Research Instrumentation Core for technical support, and members of the Ginty lab for helpful comments on the manuscript. This work was supported by NSF GRFP DG1745303 (GR), The Harvard Mahoney Neuroscience Institute Fund (AMC), The Ellen R. and Melvin J. Gordon Center for the Cure and Treatment of Paralysis (AMC), NIH grants NS119739 (AJE), NS097344 (DDG), and AT011447 (CJW and DDG), The Hock E. Tan and Lisa Yang Center for Autism Research at Harvard University (DDG), and the Edward R. and Anne G. Lefler Center for Neurodegenerative Disorders (DDG). DDG is an HHMI investigator. This article is subject to HHMI’s Open Access to Publications policy. HHMI lab heads have previously granted a nonexclusive CC BY 4.0 license to the public and a sublicensable license to HHMI in their research articles. Pursuant to those licenses, the author-accepted manuscript of this article can be made freely available under a CC BY 4.0 license immediately upon publication.

## Author Contributions

GR and DDG conceived the study. GR performed *in vivo* spinal cord electrophysiological experiments with support from AMC. GR analyzed data with assistance from AMC, AJE, and JD. GR executed SNI surgeries with assistance from ZZ and CJW. GR and DDG wrote the paper with input from all authors.

## STAR METHODS

### RESOURCE AVAILABILTY

#### Lead contact

Further information and requests for resources and reagents should be directed to and will be fulfilled by the lead contact, David Ginty.

#### Material availability

This study did not generate new unique reagents.

#### Experimental model and subject details

All animals were handled and housed in accordance with the Harvard Medical School Institutional Care and Use Committee. A mix of genetic backgrounds (C57BL/6J, CD1, 129S1/SvImJ) and female and male mice were used in this study. Animals were group housed with littermates on a 12-hour light/dark cycle. Tail biopsies and/or ear notching tissue samples were used for genotyping.

### METHOD DETAILS

#### Spared nerve injury

The spared nerve injury (SNI) model of neuropathic pain was used to induce mechanical allodynia in mice. The surgery was performed as described previously^46^. Briefly, mice were anesthetized using 2% isoflurane and an incision over the biceps femoris muscle on the lateral thigh was made to expose the sciatic nerve. The peroneal and tibial branches of the sciatic nerve were ligated and transected while sparing the sural branch. Sham surgeries involved exposure of the sciatic nerve without ligation and transection. Behavioral tests for allodynia scoring were performed at Day 0 prior to SNI surgery, Day 7 post-surgery, and Day 28 post-surgery (for animals in the chronic SNI condition).

#### *In vivo* spinal cord multielectrode array (MEA) recordings

Recordings were amplified, filtered (0.1 - 7.5 kHz bandpass), and digitized (20 kHz) using a headstage amplifier and recording controller (Intan Technologies RHD2132 and Recording Controller). Data acquisition was controlled with open-source software (Intan Technologies Recording Controller version 2.07).

*In vivo* recordings were performed on animals between 6 and 24 weeks of age. Animals were administered dexamethasone 1 to 2 hours before recording and anesthetized using urethane (1mg/kg, Sigma). Temperature of the animal was monitored and maintained (TC-344B, Warner Instruments) between 35 -37.5°C using a thermoelectric heater (C3200-6145, Honeywell) embedded in castable cement (Aremco). Surgery was performed to expose the spinal cord. An incision was made above T13 to L6 of the spine and the surrounding tissue was removed exposing the spinal column. The vertebrae between L4 and L5 were then teased apart to expose the dorsal spinal cord. The spine was then stabilized using custom clamps to prevent movement. The dura was removed from atop the spinal cord and a 32-channel silicon probe (Cambridge Neurotech ASSY-37 H4 with 200 core fiber attached-for opto-tagging) was inserted into the lateral hindpaw region of the dorsal horn.

To confirm probe placement, the hindpaw was gently brushed while monitoring multiple channels for evoked spikes. If the receptive field was not on the lateral hindpaw the probe was removed and reinserted in a new location. Recordings began twenty minutes after probe insertion. A 0.2-mm diameter, Teflon-tipped indenting probe was controlled by a dual-mode force controller (Aurora Scientific 300C-I) and used to indent the lateral hindpaw. The position, force, and displacement of the indenter were commanded with custom Matlab (version 2019a) scripts controlling a Nidaq board (National Instruments, NI USB 6259). Force steps were applied atop the minimum force required to keep the indenting probe in contact with the skin. The lateral hindpaw was stimulated with the indenting probe at a minimum of two locations which were manually determined to be receptive field hotspots for the majority of simultaneously recorded units.

#### Spike Sorting

Open-source software (JRCLUST version 3.2.5) was used to automatically sort action potentials into clusters, manually refine clusters, and classify clusters as single or multi-units^63^. The voltage traces were filtered with a differentiation filter of order 3. Frequency outliers were removed with a threshold of 10 median absolute deviations (MADs). Action potentials were detected with a threshold of 4.5 times the standard deviation of the noise. Action potentials with similar times across sites were merged and action potentials were then sorted into clusters with a density-based-clustering algorithm (clustering by fast search and find of density peaks) with cutoffs for log_10_(r) at -3 and log_10_(d) at 0.6. Clusters with a waveform correlation greater than 0.99 were automatically merged. Outlier spikes (> 6.5 MADs) were removed from each cluster.

Manual cluster curation was performed with JRCLUST split and merge tools to ensure single unit isolation. Clusters were classified as putative single units if waveforms were large with respect to baseline, a clear refractory period in the cross-correlogram (interspike intervals > 1ms) was observed, and if they were clearly distinct and separable from neighboring clusters. Spike times for single units were exported and processed in Python (3.8.5).

#### Extracellular waveform characteristics

K-means clustering was performed using waveform statistics including trough-to-peak ratio, waveform slope, and trough-to-peak duration. The somatosensory cortex (S1) waveforms and control spinal cord waveforms analyzed here were from units recorded in previously published datasets^15, 54^. K=3 was chosen to clearly separate waveforms in S1 (similar to the separation of visual cortex waveforms previously observed^55^) into two regular spiking groups and one fast-spiking group. To compare spinal cord and S1 waveforms, we used K=3 revealing less separable groups in spinal cord waveforms. No differences in extracellular waveforms were observed between sham and SNI spinal cord neurons.

#### Optical stimulation and identification of spinal cord neurons

We used the optical tagging strategy described previously^15^ to identify genetically defined dorsal horn interneuron populations. Briefly, interneurons expressing excitatory opsins were opto-tagged by delivering pulses (1-20ms) of blue light (4-10 mW/mm^2^ at fiber tip) to the surface of the spinal cord through an optical fiber (200µm core diameter; NA = 0.66) attached to Cambridge Neurotech ASSY-37 H4 optrodes. Light was delivered from a 470 nm LED (M470F3, Thorlabs). Optical stimulation was performed after mechanical stimulation. At the conclusion of each experiment, 25µl of 5mM NBQX (5 mM, Tocris, dissolved in H_2_O) was applied to the surface of the spinal cord to block possible recurrent glutamatergic transmission. Abolished tactile responses to steps of indentation on the hindpaw determined efficient block of glutamatergic transmission (normally between 10 and 20 minutes after NBQX application, Figures S3A and S3B) and optical stimulation was repeated. Neurons that responded to stimulation both before and after NBQX application are determined to be opto-tagged. A modified stimulus-associated spike latency test (SALT ^54, 64^) was additionally used to confirm short light-evoked spike latencies (< 10ms, Figure S3C) and low spike jitter in opto-tagged units.

#### Indentation and brush response properties

Tactile responsive single units were identified by responding to 500ms steps of indentation at varying innocuous forces, between 1 and 75mN. We subdivided step indentations into 3 different time periods to monitor different aspects of neuronal responses: ON response: 0-50ms after stimulus onset; OFF response: 0-50ms after stimulus offset; and Sustained response: 0-200ms before stimulus offset. Thresholds for all units were determined by bootstrapping the baseline firing rate 1000 times to generate 95% confidence intervals and detecting the smallest stimulus within the ON/OFF/Sustained response windows that exceeds the upper bound.

Units that had no response threshold (only baseline firing detected) were excluded. Baseline firing was computed over a 1.5s period prior to the indentation stimuli. Peristimulus time histograms (PSTHs) were generated to show the average response across all units within a condition, genotype, and/or that have been optically tagged. These PSTHs were created with 10ms time bins unless otherwise noted.

The lateral hindpaw was lightly stroked with a soft 1.2mm wide brush for nine minutes (the brush and amount of time stroking the paw are consistent with the behavioral dynamic brush assay). For each neuron responding above baseline to brush stimuli, maximum evoked firing rates were computed for each minute of brush stimulation and these values were then averaged across all brush sessions.

#### Firing correlation analyses

Signal, noise, and synchrony cross-correlations were calculated between pairs of simultaneously recorded neurons. To calculate signal correlations, spiking responses to various stimuli were averaged across trials and the Pearson correlation coefficient of mean responses (PSTHs, 50ms bins) between pairs of neurons were computed. Trial-by-trial spike count correlations were used to determine noise correlations. Cross-correlograms were generated (using Python module Scipy.Signal) from spiking data in 1ms bins to determine paired firing synchrony at 0 time lag.

Population coupling was calculated as previously described^47^. Briefly, simultaneously recorded single unit activity was summed into a population rate with 1ms resolution. The population rate was used to compute a spike-triggered population rate for each unit (not including the spikes of that unit). To compare between recordings, conditions, and genotypes each spike-triggered population rate was normalized by subtracting the median spike-triggered population rate of shuffled spiking data (randomized spike times) for each experiment. These values reflect normalized synchronous firing at a population level and are plotted as normalized population firing rates.

#### Indentation latencies and jitter measurements

Latency and jitter of single units were calculated in response to 10mN and 75mN step indentations of the skin. The distribution of first spike latencies to each force step was compared to a shuffled distribution (shuffled at least 100 times). The time when the distribution exceeded the 95% confidence interval of the shuffled distribution was determined to be the latency. The standard deviation of the first spike latencies across trials was then calculated to determine the jitter. A minimum of 50 trials were used to calculate latencies and jitter.

#### Spinal cord immunohistochemistry of free-floating sections

Adult mice were anesthetized with isoflurane and perfused with 10mL of 1X Phosphate Buffer Saline (PBS), followed by 20 mL of 4% paraformaldehyde (PFA) in PBS at room temperature. Vertebral columns were dissected and were post-fixed in 4% PFA at 4°C for 24 hr. Lumbar spinal cord coronal sections (60 µm) were cut on a vibrating blade microtome (Leica VT100S) and processed for immunohistochemistry as described previously^19, 58^. Briefly, tissue samples were rinsed in 50% ethanol/water solution for 30 min to allow for enhanced antibody penetration followed by three washes in high salt PBS each lasting 10 min. The tissue was then incubated with primary antibodies (goat anti-mCherry (1:1000, AB0040, Scigen), rabbit anti-GFP (1:1000, A-11122, Thermo Fisher Scientific)) in high salt PBS containing 0.3% Triton X-100 (HS PBSt) for 48 hr at 4°C. The tissue was washed in HS PBSt, then incubated in a secondary antibody solution in HS PBSt overnight at 4°C. Secondary antibodies included species-specific Alexa Fluor 488 and 546 conjugated IgGs (1:500; Life Technologies), and IB4 (1:500; Alexa 647 conjugated, L21411, Molecular Probes). Tissue sections were then mounted on glass slides, coverslipped with Fluoromount Aqueous Mounting Medium (Sigma) and stored at 4°C.

#### Dynamic brush assay

Dynamic mechanical allodynia was determined as previously described^34^. Briefly, hypersensitivity was measured by stroking the lateral side of the injured or sham hindpaw from heel to toe with a soft 1.2mm wide brush. Behaviors were scored from 0 to 3. No movement or a very fast lifting of the stimulated paw for less than 1 sec scored as a 0. After nerve injury several pain-suggestive responses can be observed, such as sustained lifting (2 sec or more) of the stimulated paw (scored as 1); lateral kicking/ flinching of stimulated hindpaw (scored as 2); and licking of the stimulated paw (scored as 3). Stroking was performed for three-minute periods and repeated three times. The highest score per period was then averaged for each mouse. Sham mice scored an average score of ∼0 seven days post SNI surgery and SNI mice scored ∼2.3. Efficient induction of mechanical allodynia was determined by a score >1.5.

#### Quantification and statistical analysis

Statistical tests were conducted using the SciPy stats module (Python 3.8.5) or GraphPad Prism. Both non-parametric tests and parametric tests were used, depending on data normality, for comparing two independent groups (Mann-Whitney U test or unpaired t test), and multiple groups (Kruskal-Wallis test/ one-way ANOVA or two-way ANOVA for multiple groups with multiple timepoints). All post-hoc comparisons performed are indicated in the figure legends. Only significantly different p-values are reported and a p < 0.05 was considered significant. All error bars plotted display 95% confidence intervals (CI) unless otherwise noted. All box and whisker plots show median, lower and upper quartiles, and minimum to maximum values. Additional details on sample sizes and statistical tests for each experiment can be found in the figure legends, main text, and supplemental table.

#### Data and code availability

All data reported in this study will be shared by the lead contact upon request. Code is also available upon request.

---

**Figure S1.**
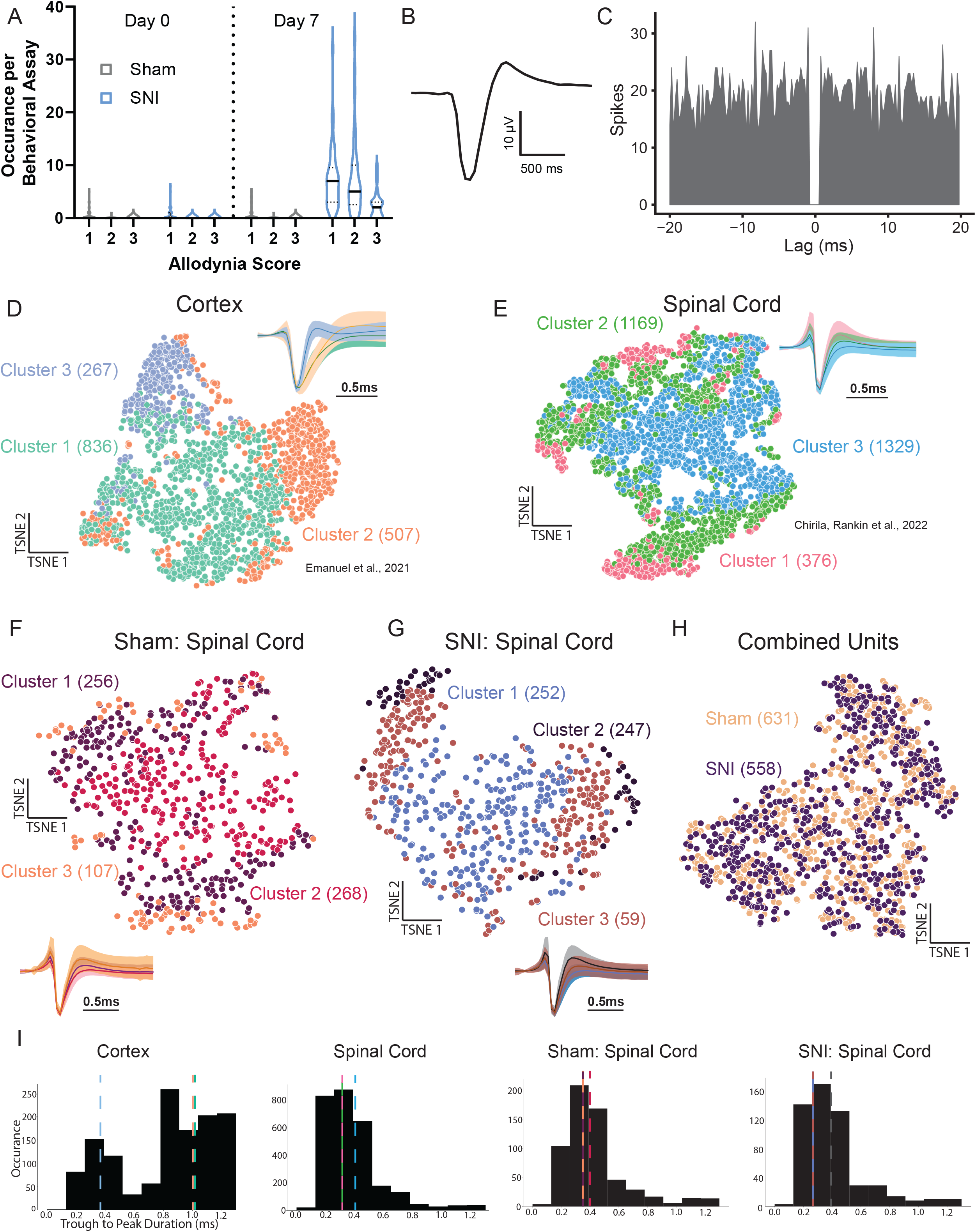
Allodynic behaviors and extracellular waveform features of spinal cord interneurons. **A.** Behaviors exhibited over dynamic brush assay per allodynia score category in sham (N=22) and SNI (N=19) mice. 1: sustained lifting (2 sec or more) of the stimulated paw. 2: lateral kicking/ flinching of stimulated hindpaw. 3: licking of the stimulated paw. **B.** Example extracellular waveform from a DH neuron. **C.** Corresponding auto-correlogram for the neuron shown in Figure S1B. **D.** t-SNE visualization of somatosensory cortex extracellular waveforms clustered using k-means clustering (n= 1610 units; see Methods). As previously reported^55^, three primary waveform subtypes emerged: two regular-spiking (green, cluster 1; orange, cluster 2) and one fast-spiking (light blue, cluster 3). Normalized mean waveforms in different clusters are shown ±SD. **E.** Extracellular waveforms for DH units (n = 2874 units) clustered as in (D). Unlike cortex, DH waveforms are not clearly separated into three groups (pink, cluster 1; green, cluster 2; blue, cluster 3). **F.** Sham DH unit extracellular waveforms (n = 631 units) clustered as in D (dark purple, cluster 1; red, cluster 2; orange, cluster 3). **G.** SNI DH unit extracellular waveforms (n = 558 units) clustered as in D (blue, cluster 1; black, cluster 2; brown, cluster 3). **H.** Combined sham and SNI unit waveforms do not cluster separately. **I.** Trough-to-peak durations for each region/condition. Cortical unit waveforms have a bimodal distribution, highlighting the duration difference between fast-spiking (dashed purple line, cluster 3; median: 0.35ms) and regular-spiking (dashed green line, cluster 1; median: 1.05ms; dashed orange line, cluster 2; median: 1.00ms) units. Spinal cord unit waveforms have a unimodal distribution (medians for cluster per cluster plotted as dashed lines).

**Figure S2.**
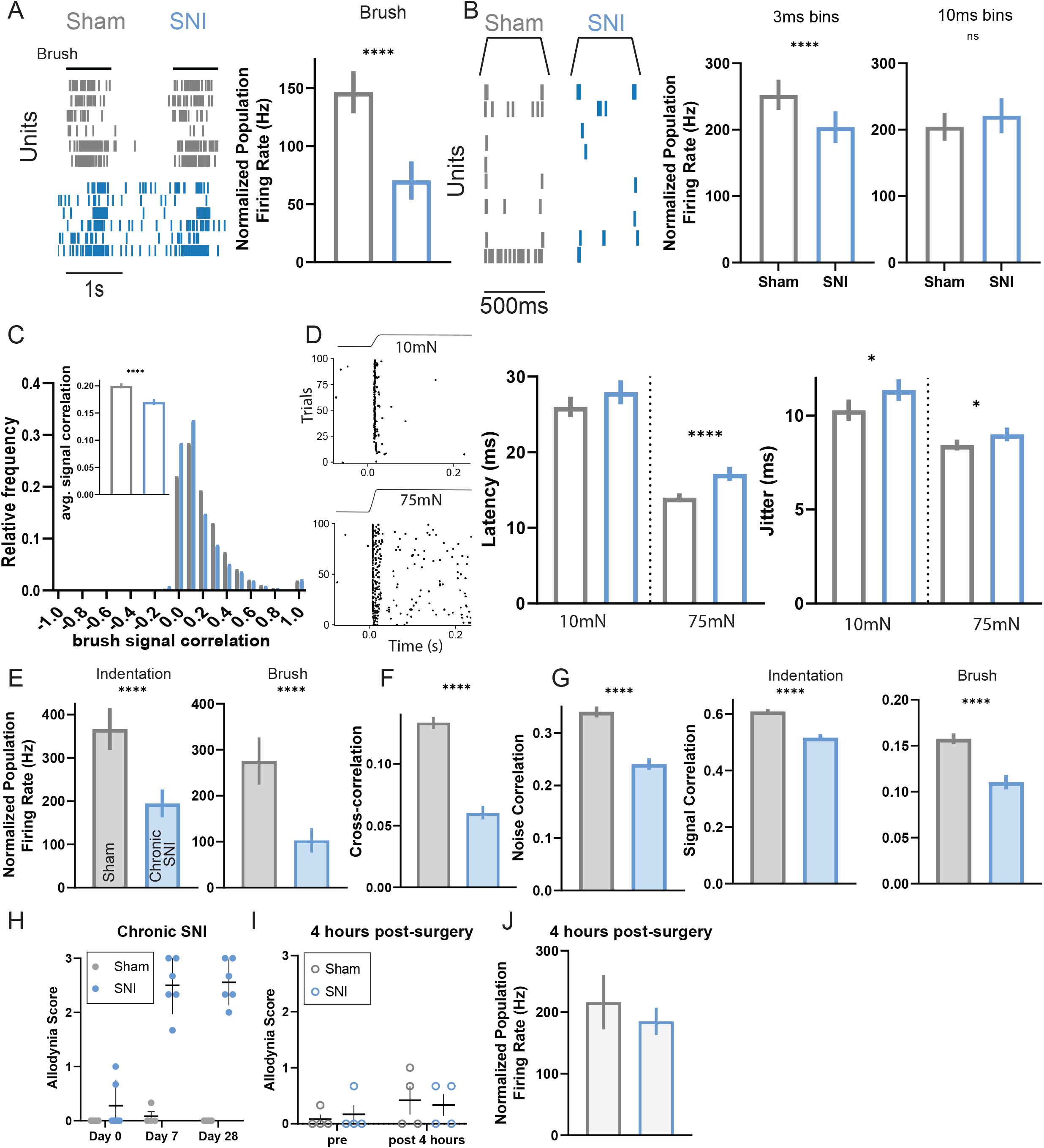
Temporal organization of DH firing across milliseconds and for hours to weeks after SNI surgery. **A.** Raster plots (left) showing temporal alignment of simultaneously recorded units responding to skin stroke in sham and SNI conditions followed by population coupling quantified as normalized population firing rate (right). Mann-Whitney *U* test. **B.** Raster plots showing temporal alignment of simultaneously recorded units responding to 50mN indentation steps (left), population coupling with expanded 3ms time bins (middle) and population coupling with expanded 10ms time bins (right). Mann-Whitney *U* tests. **C.** Distribution of brush signal correlations for pairs of DH neurons. Insets: average signal correlations. Mann-Whitney *U* test. **D.** Left: raster plots of one sham neuron responding to the onset of 10mN and 75mN indentation steps across trials. Right: quantification of latency and jitter to indentation steps across neurons. **E.** Population coupling to both indentation (left) and brush (right) stimuli after chronic SNI (28 - 35 days post-surgery). Mann-Whitney *U* test. **F.** Synchrony cross-correlations for neuron pairs after chronic SNI. Mann-Whitney *U* test. **G.** Noise (left) and signal correlations (indentation: middle, brush: right) for neuron pairs after chronic SNI. Mann-Whitney *U* tests. **H.** Dynamic allodynia score compared at Day 0 (prior to surgery), Day 7 (post-surgery), and Day 28 (post-surgery) between sham (N=4) and SNI (N=6) mice. One-way ANOVA with post-hoc Tukey’s test [F (5, 24) = 64.28; p < 0.0001]. SNI day 7 and day 28 were significantly different from all other times points and conditions (****). Error bars: SEM **I.** Dynamic allodynia score compared prior to surgery and 4 hours post-surgery in sham (N=4) and SNI (N=4) mice. Error bars: SEM **J.** Population coupling 4 hours post nerve injury. Mann-Whitney *U* test. Bars: mean. Error bars: 95% CI. Number of animals/ cells (N). *p < 0.05, **p < 0.01, ***p < 0.001, ****p < 0.0001. See Table S1 for experimental details.

**Figure S3.**
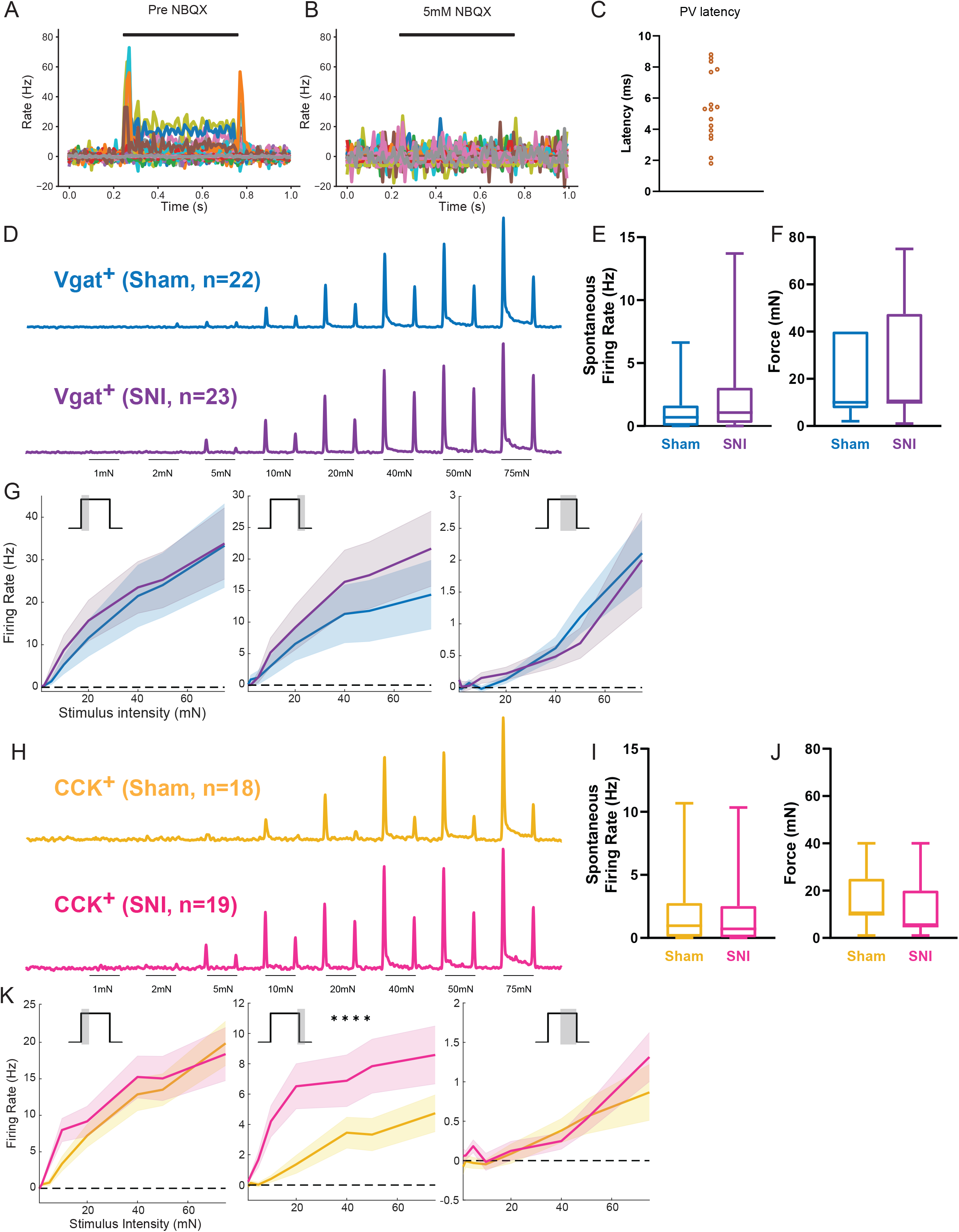
Characterizing Vgat^+^ and CCK^+^ interneuron activity following SNI. **A.** Simultaneously recorded units responding to 50mN steps of indentation (black bar). **B.** Indentation responses to 50mN step indentations (black bar) abolished by glutamatergic block using NBQX applied to the surface of the spinal cord. **C.** Average latency to first spike after optical stimulation for PV^+^ DH interneurons in the sham condition. **D.** Mean baseline-subtracted firing rate PSTHs for sham and SNI Vgat^+^ neurons. **E.** Indentation thresholds across Vgat^+^ neurons in sham (N=4, n=22) and SNI (N=5, n=23) mice. **F.** Spontaneous firing rates of Vgat^+^ interneurons. **G.** Average baseline-subtracted firing rates (±SEM) for Vgat^+^ neurons in sham and SNI groups at step indentation On (left), Off (middle), and Sustained (right) periods. **H.** Mean baseline-subtracted firing rate PSTHs for sham and SNI CCK^+^ neurons. **I.** Indentation thresholds across CCK^+^ neurons in Sham (N=4, n=18) and SNI (N=6, n=19) mice. **J.** Spontaneous firing rates of CCK^+^ interneurons. **K.** Average baseline-subtracted firing rates (±SEM) for CCK^+^ neurons in sham and SNI groups at step indentation On, Off, and Sustained periods. Off: two-way ANOVA [F (1,280) = 29.77, p < 0.0001]. Bars: mean. Error bars: 95% CI. Number of animals/ cells (N/n). *p < 0.05, **p < 0.01, ***p < 0.001.

**Figure S4.**
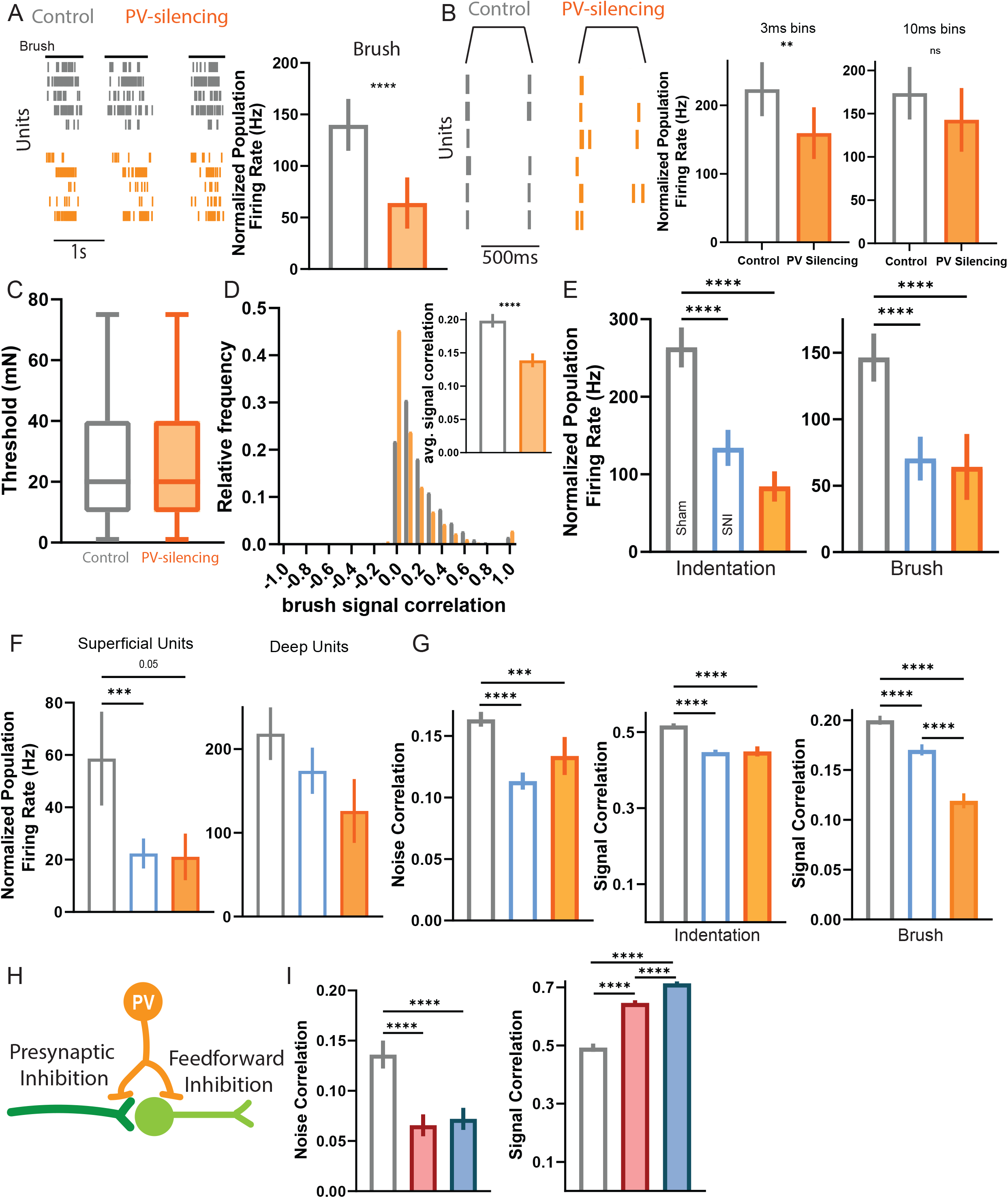
Altered correlated activity is comparable between PV-silencing and SNI conditions. **A.** Raster plots showing temporal alignment of simultaneously recorded units responding to strokes (right) in sham and SNI conditions followed by population coupling quantified as normalized population firing rate. Mann-Whitney *U* test. **B.** Raster plots showing temporal alignment of simultaneously recorded units responding to 50mN indentation steps (left), population coupling with expanded 3ms time bins (middle) and population coupling with expanded 10ms time bins (right). Mann-Whitney U tests. **C.** Indentation thresholds between control (N=4) and PV-silencing mice (N=4). Mann-Whitney *U* test. **D.** Distribution of brush signal correlations for pairs of DH neurons. Insets: average signal correlations. Mann-Whitney *U* test. **E.** Population coupling to both indentation (left) and brush (right) stimuli comparing a randomly shuffled subset of the sham and SNI datasets with the PV-silencing dataset. Indentation: One-way ANOVA with post-hoc Tukey’s test [F (2, 353) = 18.91; p < 0.0001]. Brush: One-way ANOVA with post-hoc Tukey’s test [F (2, 371) = 12.39; p < 0.0001]. **F.** Superficial (left) and deep (right) population coupling comparing a shuffled subset of the sham and SNI datasets with the PV-silencing dataset. Superficial: One-way ANOVA with post-hoc Tukey’s test [F (2, 81) = 6.703; p = 0.0020]. **G.** Noise (left) and signal correlations (indentation: middle, brush: right) for neuron pairs from a randomly shuffled subset of the sham and SNI datasets with the PV-silencing dataset. Noise: One-way ANOVA with post-hoc Tukey’s test [F (2, 4184) = 8.124; p = 0.0003]. Signal (indentation): One-way ANOVA with post-hoc Tukey’s test [F (2, 4184) = 8.124; p = 0.0003]. **H.** Diagram of presynaptic and feedforward inhibitory synapses formed by PV^+^ DH neurons. **I.** Noise and signal correlations (indentation) for neuron pairs. Noise: Kruskal-Wallis H test with post-hoc Dunn’s test (H[2,6099] = 60.27; p < 0.0001). Signal: Kruskal-Wallis H test with post-hoc Dunn’s test (H[2, 6099] = 618.2; p < 0.0001). Bars: mean. Error bars: 95% CI. Number of animals/ cells (N). *p < 0.05, **p < 0.01, ***p < 0.001. See Table S1 for experimental details.

## References

1. von Hehn, C.A., Baron, R., and Woolf, C.J. (2012). Deconstructing the Neuropathic Pain Phenotype to Reveal Neural Mechanisms. Neuron 73, 638–652. 10.1016/J.NEURON.2012.02.008.

2. Bouhassira, D., Attal, N., Alchaar, H., Boureau, F., Brochet, B., Bruxelle, J., Cunin, G., Fermanian, J., Ginies, P., Grun-Overdyking, A., et al. (2005). Comparison of pain syndromes associated with nervous or somatic lesions and development of a new neuropathic pain diagnostic questionnaire (DN4). Pain 114, 29–36. 10.1016/J.PAIN.2004.12.010.

3. Goldberg, D.S., and McGee, S.J. (2011). Pain as a global public health priority. BMC Public Health 11, 770. 10.1186/1471-2458-11-770.

4. Munro, G., Ahring, P.K., and Mirza, N.R. (2009). Developing analgesics by enhancing spinal inhibition after injury: GABAA receptor subtypes as novel targets. Trends Pharmacol Sci 30, 453–459. 10.1016/J.TIPS.2009.06.004.

5. Dworkin, R.H., Backonja, M., Rowbotham, M.C., Allen, R.R., Argoff, C.R., Bennett, G.J., Bushnell, M.C., Farrar, J.T., Galer, B.S., Haythornthwaite, J.A., et al. (2003). Advances in Neuropathic Pain: Diagnosis, Mechanisms, and Treatment Recommendations. Arch Neurol 60, 1524–1534. 10.1001/ARCHNEUR.60.11.1524.

6. Baron, R. (2006). Mechanisms of Disease: neuropathic pain—a clinical perspective. Nat Clin Pract Neurol 2, 95–106. 10.1038/ncpneuro0113.

7. Jensen, M.P., Chodroff, M.J., and Dworkin, R.H. (2007). The impact of neuropathic pain on health-related quality of life: review and implications. Neurology 68, 1178–1182. 10.1212/01.wnl.0000259085.61898.9e.

8. Woolf, C.J., and Mannion, R.J. (1999). Neuropathic pain: aetiology, symptoms, mechanisms, and management. The Lancet 353, 1959–1964. 10.1016/S0140-6736(99)01307-0.

9. Moehring, F., Halder, P., Seal, R.P., and Stucky, C.L. (2018). Uncovering the Cells and Circuits of Touch in Normal and Pathological Settings. Neuron 100, 349–360. 10.1016/J.NEURON.2018.10.019.

10. Kim, W., Kim, S.K., and Nabekura, J. (2017). Functional and structural plasticity in the primary somatosensory cortex associated with chronic pain. J Neurochem 141, 499–506. 10.1111/JNC.14012.

11. Bliss, T.V.P., Collingridge, G.L., Kaang, B.K., and Zhuo, M. (2016). Synaptic plasticity in the anterior cingulate cortex in acute and chronic pain. Nature Reviews Neuroscience 2016 17:8 17, 485–496. 10.1038/NRN.2016.68.

12. Latremoliere, A., and Woolf, C.J. (2009). Central Sensitization: A Generator of Pain Hypersensitivity by Central Neural Plasticity. J Pain 10, 895–926. 10.1016/J.JPAIN.2009.06.012.

13. Melzack, R., and Wall, P.D. (1965). Pain mechanisms: a new theory. Science 150, 971– 979.

14. Woolf, C.J. (1983). Evidence for a central component of post-injury pain hypersensitivity. Nature 306, 686–688. 10.1038/306686a0.

15. Chirila, A.M., Rankin, G., Tseng, S.-Y., Emanuel, A.J., Chavez-Martinez, C.L., Zhang, D., Harvey, C.D., and Ginty, D.D. (2022). Mechanoreceptor signal convergence and transformation in the dorsal horn flexibly shape a diversity of outputs to the brain. Cell 185, 4541–4559.e23. 10.1016/J.CELL.2022.10.012.

16. Handler, A., and Ginty, D.D. (2021). The mechanosensory neurons of touch and their mechanisms of activation. Nat Rev Neurosci 22, 521–537. 10.1038/s41583-021-00489-x.

17. Paixão, S., Loschek, L., Gaitanos, L., Alcalà Morales, P., Goulding, M., and Klein, R. (2019). Identification of Spinal Neurons Contributing to the Dorsal Column Projection Mediating Fine Touch and Corrective Motor Movements. Neuron 104, 749–764.e6. 10.1016/j.neuron.2019.08.029.

18. Choi, S., Hachisuka, J., Brett, M.A., Magee, A.R., Omori, Y., Iqbal, N. ul A., Zhang, D., DeLisle, M.M., Wolfson, R.L., Bai, L., et al. (2020). Parallel ascending spinal pathways for affective touch and pain. Nature 587, 258. 10.1038/S41586-020-2860-1.

19. Abraira, V.E., Kuehn, E.D., Chirila, A.M., Heintz, N., Hughes, D.I., and Ginty, D.D. (2017). The Cellular and Synaptic Architecture of the Mechanosensory Dorsal Horn. Cell 168, 295–310. 10.1016/j.cell.2016.12.010.

20. Peirs, C., Dallel, R., and Todd, A.J. (2020). Recent advances in our understanding of the organization of dorsal horn neuron populations and their contribution to cutaneous mechanical allodynia. J Neural Transm 127, 505. 10.1007/S00702-020-02159-1.

21. Hughes, D.I., and Todd, A.J. (2020). Central Nervous System Targets: Inhibitory Interneurons in the Spinal Cord. Neurotherapeutics 17, 874–885. 10.1007/S13311-020-00936-0/FIGURES/4.

22. Eccles, J.C., Schmidt, R., and Willis, W.D. (1963). PHARMACOLOGICAL STUDIES ON PRESYNAPTIC INHIBITION. J Physiol 168, 500–530.

23. Eccles, J.C., Schmidt, R.F., and Willis, W.D. (1963). DEPOLARIZATION OF THE CENTRAL TERMINALS OF CUTANEOUS AFFERENT FIBERS. J Neurophysiol 26, 646–661. 10.1152/jn.1963.26.4.646.

24. Koch, S.C., del Barrio, M.G., Dalet, A., Gatto, G., Günther, T., Zhang, J., Seidler, B., Saur, D., Schüle, R., and Goulding, M. (2017). RORβ Spinal Interneurons Gate Sensory Transmission during Locomotion to Secure a Fluid Walking Gait. Neuron 96, 1419–1431.e5. 10.1016/j.neuron.2017.11.011.

25. Orefice, L.L., Zimmerman, A.L., Chirila, A.M., Sleboda, S.J., Head, J.P., and Ginty, D.D. (2016). Peripheral Mechanosensory Neuron Dysfunction Underlies Tactile and Behavioral Deficits in Mouse Models of ASDs. Cell 166, 299–313. 10.1016/j.cell.2016.05.033.

26. Zimmerman, A.L., Kovatsis, E.M., Pozsgai, R.Y., Tasnim, A., Zhang, Q., and Ginty, D.D. (2019). Distinct Modes of Presynaptic Inhibition of Cutaneous Afferents and Their Functions in Behavior. Neuron 102, 420–434.e8. 10.1016/j.neuron.2019.02.002.

27. Lever, I., Cunningham, J., Grist, J., Yip, P.K., and Malcangio, M. (2003). Release of BDNF and GABA in the dorsal horn of neuropathic rats. European Journal of Neuroscience 18, 1169–1174. 10.1046/J.1460-9568.2003.02848.X.

28. Moore, K.A., Kohno, T., Karchewski, L.A., Scholz, J., Baba, H., and Woolf, C.J. (2002). Partial Peripheral Nerve Injury Promotes a Selective Loss of GABAergic Inhibition in the Superficial Dorsal Horn of the Spinal Cord. Journal of Neuroscience 22, 6724–6731. 10.1523/JNEUROSCI.22-15-06724.2002.

29. Castro-Lopes, J.M., Tavares, I., and Coimbra, A. (1993). GABA decreases in the spinal cord dorsal horn after peripheral neurectomy. Brain Res 620, 287–291. 10.1016/0006-8993(93)90167-L.

30. Torsney, C., and MacDermott, A.B. (2006). Disinhibition opens the gate to pathological pain signaling in superficial neurokinin 1 receptor-expressing neurons in rat spinal cord. J Neurosci 26, 1833–1843. 10.1523/JNEUROSCI.4584-05.2006.

31. Boyle, K.A., Gradwell, M.A., Yasaka, T., Callister, R.J., Graham, B.A., and Hughes Correspondence, D.I. (2019). Defining a Spinal Microcircuit that Gates Myelinated Afferent Input: Implications for Tactile Allodynia. Cell Rep 28, 526–540. 10.1016/j.celrep.2019.06.040.

32. Cui, L., Miao, X., Liang, L., Abdus-Saboor, I., Olson, W., Fleming, M.S., Ma, M., Tao, Y.-X., and Luo, W. (2016). Identification of Early RET+ Deep Dorsal Spinal Cord Interneurons in Gating Pain. Neuron 91, 1137–1153. 10.1016/J.NEURON.2016.07.038.

33. Lu, Y., Dong, H., Gao, Y., Gong, Y., Ren, Y., Gu, N., Zhou, S., Xia, N., Sun, Y.Y., Ji, R.R., et al. (2013). A feed-forward spinal cord glycinergic neural circuit gates mechanical allodynia. J Clin Invest 123, 4050. 10.1172/JCI70026.

34. Duan, B., Cheng, L., Bourane, S., Britz, O., Padilla, C., Garcia-Campmany, L., Krashes, M., Knowlton, W., Velasquez, T., Ren, X., et al. (2014). Identification of Spinal Circuits Transmitting and Gating Mechanical Pain. Cell 159, 1417–1432. 10.1016/J.CELL.2014.11.003.

35. Foster, E., Wildner, H., Tudeau, L., Haueter, S., Ralvenius, W.T., Jegen, M., Johannssen, H., Hösli, L., Haenraets, K., Ghanem, A., et al. (2015). Targeted Ablation, Silencing, and Activation Establish Glycinergic Dorsal Horn Neurons as Key Components of a Spinal Gate for Pain and Itch. Neuron 85, 1289–1304. 10.1016/J.NEURON.2015.02.028.

36. Petitjean, H., Pawlowski, S.A., Fraine, S.L., Sharif, B., Hamad, D., Fatima, T., Berg, J., Brown, C.M., Jan, L.Y., Ribeiro-da-Silva, A., et al. (2015). Dorsal Horn Parvalbumin Neurons Are Gate-Keepers of Touch-Evoked Pain after Nerve Injury. Cell Rep 13, 1246– 1257. 10.1016/J.CELREP.2015.09.080.

37. Cardin, J.A. (2018). Inhibitory Interneurons Regulate Temporal Precision and Correlations in Cortical Circuits. Trends Neurosci 41, 689–700. 10.1016/J.TINS.2018.07.015.

38. Veit, J., Hakim, R., Jadi, M.P., Sejnowski, T.J., and Adesnik, H. (2017). Cortical gamma band synchronization through somatostatin interneurons. Nature Neuroscience 2017 20:7 20, 951–959. 10.1038/NN.4562.

39. Cardin, J.A., Carlén, M., Meletis, K., Knoblich, U., Zhang, F., Deisseroth, K., Tsai, L.H., and Moore, C.I. (2009). Driving fast-spiking cells induces gamma rhythm and controls sensory responses. Nature 2009 459:7247 459, 663–667. 10.1038/NATURE08002.

40. Buia, C., and Tiesinga, P. (2006). Attentional modulation of firing rate and synchrony in a model cortical network. J Comput Neurosci 20, 247–264. 10.1007/S10827-006-6358-0/METRICS.

41. English, D.F., McKenzie, S., Evans, T., Kim, K., Yoon, E., and Buzsáki, G. (2017). Pyramidal Cell-Interneuron Circuit Architecture and Dynamics in Hippocampal Networks. Neuron 96, 505–520.e7. 10.1016/J.NEURON.2017.09.033.

42. Batista-Brito, R., Vinck, M., Ferguson, K.A., Chang, J.T., Laubender, D., Lur, G., Mossner, J.M., Hernandez, V.G., Ramakrishnan, C., Deisseroth, K., et al. (2017). Developmental Dysfunction of VIP Interneurons Impairs Cortical Circuits. Neuron 95, 884–895.e9. 10.1016/J.NEURON.2017.07.034.

43. Zeilhofer, H.U., Wildner, H., and Yévenes, G.E. (2012). Fast Synaptic Inhibition in Spinal Sensory Processing and Pain Control. Physiol Rev 92, 193–235. 10.1152/physrev.00043.2010.

44. Chen, J.T., Guo, D., Campanelli, D., Frattini, F., Mayer, F., Zhou, L., Kuner, R., Heppenstall, P.A., Knipper, M., and Hu, J. (2014). Presynaptic GABAergic inhibition regulated by BDNF contributes to neuropathic pain induction. Nat Commun 5, 5331. 10.1038/ncomms6331.

45. Malan, T.P., Mata, H.P., and Porreca, F. (2002). Spinal GABAAand GABABReceptor Pharmacology in a Rat Model of Neuropathic Pain. Anesthesiology 96, 1161–1167. 10.1097/00000542-200205000-00020.

46. Decosterd, I., and Woolf, C.J. (2000). Spared nerve injury: an animal model of persistent peripheral neuropathic pain. Pain 87, 149–158. 10.1016/S0304-3959(00)00276-1.

47. Okun, M., Steinmetz, N.A., Cossell, L., Iacaruso, M.F., Ko, H., Barthó, P., Moore, T., Hofer, S.B., Mrsic-Flogel, T.D., Carandini, M., et al. (2015). Diverse coupling of neurons to populations in sensory cortex. Nature 521, 511–515. 10.1038/nature14273.

48. Sweeney, Y., and Clopath, C. (2020). Population coupling predicts the plasticity of stimulus responses in cortical circuits. Elife 9. 10.7554/eLife.56053.

49. Woolf, C.J., and Fitzgerald, M. (1986). Somatotopic organization of cutaneous afferent terminals and dorsal horn neuronal receptive fields in the superficial and deep laminae of the rat lumbar spinal cord. Journal of Comparative Neurology 251, 517–531. 10.1002/CNE.902510407.

50. Li, L., Rutlin, M., Abraira, V.E., Cassidy, C., Kus, L., Gong, S., Jankowski, M.P., Luo, W., Heintz, N., Koerber, H.R., et al. (2011). The functional organization of cutaneous low-threshold mechanosensory neurons. Cell 147, 1615–1627. 10.1016/j.cell.2011.11.027.

51. Koerber, H.R., and Brown, P.B. (1982). Somatotopic organization of hindlimb cutaneous nerve projections to cat dorsal horn. J Neurophysiol 48, 481–489. 10.1152/JN.1982.48.2.481.

52. Cohen, M.R., and Kohn, A. (2011). Measuring and interpreting neuronal correlations. Nature Neuroscience 2011 14:7 14, 811–819. 10.1038/nn.2842.

53. Niell, C.M., and Stryker, M.P. (2008). Highly Selective Receptive Fields in Mouse Visual Cortex. Journal of Neuroscience 28, 7520–7536. 10.1523/JNEUROSCI.0623-08.2008.

54. Emanuel, A.J., Lehnert, B.P., Panzeri, S., Harvey, C.D., and Ginty, D.D. (2021). Cortical responses to touch reflect subcortical integration of LTMR signals. Nature 600, 680–685. 10.1038/s41586-021-04094-x.

55. Xiaoxuan Jia, X., Siegle, J.H., Bennett, C., Gale, S.D., Denman, D.J., Koch, C., and Olsen, S.R. (2019). High-density extracellular probes reveal dendritic backpropagation and facilitate neuron classification. J Neurophysiol 121, 1831–1847. 10.1152/jn.00680.

56. Cao, B., Scherrer, G., and Chen, L. (2022). Spinal cord retinoic acid receptor signaling gates mechanical hypersensitivity in neuropathic pain. Neuron 110, 4108–4124.e6. 10.1016/J.NEURON.2022.09.027.

57. Liu, Y., Latremoliere, A., Li, X., Zhang, Z., Chen, M., Wang, X., Fang, C., Zhu, J., Alexandre, C., Gao, Z., et al. (2018). Touch and tactile neuropathic pain sensitivity are set by corticospinal projections. Nature 561, 547–550. 10.1038/s41586-018-0515-2.

58. Hughes, D.I., Sikander, S., Kinnon, C.M., Boyle, K.A., Watanabe, M., Callister, R.J., and Graham, B.A. (2012). Morphological, neurochemical and electrophysiological features of parvalbumin-expressing cells: a likely source of axo-axonic inputs in the mouse spinal dorsal horn. J Physiol 590, 3927–3951. 10.1113/jphysiol.2012.235655.

59. Orefice, L.L., Mosko, J.R., Morency, D.T., Wells, M.F., Tasnim, A., Mozeika, S.M., Ye, M., Chirila, A.M., Emanuel, A.J., Rankin, G., et al. (2019). Targeting Peripheral Somatosensory Neurons to Improve Tactile-Related Phenotypes in ASD Models. Cell 178, 867–886.e24. 10.1016/j.cell.2019.07.024.

60. Rao, R.P.N., and Ballard, D.H. (1999). Predictive coding in the visual cortex: a functional interpretation of some extra-classical receptive-field effects. Nature Neuroscience 1999 2:1 2, 79–87. 10.1038/4580.

61. Brown, A.G., and Fyffe, R.E. (1981). Form and function of dorsal horn neurones with axons ascending the dorsal columns in cat. J Physiol 321, 31–47. 10.1113/jphysiol.1981.sp013970.

62. Brown, A.G., Brown, P.B., Fyffe, R.E., and Pubols, L.M. (1983). Receptive field organization and response properties of spinal neurones with axons ascending the dorsal columns in the cat. J Physiol 337, 575–588. 10.1113/jphysiol.1983.sp014643.

63. Jun, J.J., Mitelut, C., Lai, C., Gratiy, S., Anastassiou, C., and Harris, T.D. (2017). Real-time spike sorting platform for high-density extracellular probes with ground-truth validation and drift correction. bioRxiv, 101030. 10.1101/101030.

64. Kvitsiani, D., Ranade, S., Hangya, B., Taniguchi, H., Huang, J.Z., and Kepecs, A. (2013). Distinct behavioural and network correlates of two interneuron types in prefrontal cortex. Nature 498, 363. 10.1038/NATURE12176.

