## supplemental table 1 for "Nerve injury disrupts temporal processing in the spinal cord dorsal horn through alterations in PV^+^ interneurons"

**Table S1**

N=animals n=pairs of cells

| Figure 2 | panel | sub-panel | Sham (N/n) | SNI (N/n) |
|---|---|---|---|---|
| # | **A** | indentation | 22/642 | 19/479 |
| # | **B** | superficial | 21/120 | 18/102 |
| | | deep | 21/485 | 18/364 |
| | **C** | | 21/9351 | 19/7535 |
| | **D** | | 22/9921 | 17/7008 |
| | **E** | | 21/9351 | 19/7535 |

| Figure S2 | panel | sub-panel | Sham (N/n) | SNI (N/n) |
|---|---|---|---|---|
| # | **A** | | 15/318 | 12/321 |
| # | **B** | 3ms | 22/642 | 19/479 |
| | | 10ms | 22/642 | 19/479 |
| # | **C** | | 15/5040 | 12 /4840 |
| # | **D** | 10mN | | |
| | | latency | 21/179 | 18/151 |
| | | jitter | 21/179 | 18/151 |
| | | 75mN | | |
| | | latency | 21/447 | 18/318 |
| | | jitter | 21/447 | 18/318 |
| # | **E** | indentation | 4/103 | 5/116 |
| | | brush | 4/152 | 5/143 |
| | **F** | | 4/ 2959 | 5 /2041 |
| | **G** | noise | 4/ 2959 | 5 /2041 |
| | | indentation | 4/ 2959 | 5 /2041 |
| | | brush | 4 /4281 | 5/ 2296 |
| | **I** | | 4 | 4 |
| | **J** | | 4/120 | 4/ 87 |

### n= number of cells

| Figure 4 | panel | sub-panel | Control (N/n) | Condition (N/n) | Condition (N/n) |
|---|---|---|---|---|---|
| | | | | **PV-silencing** | |
| # | **A** | indentation | 4/ 99 | 4/108 | |
| # | **B** | superficial | 4/ 21 | 4 /24 | |
| | | deep | 4/ 78 | 4 /84 | |
| | **C** | | 4/1410 | 4/1384 | |
| | **D** | noise | 4/1410 | 4/1384 | |
| | | indentation | 4/1410 | 4/1384 | |
| | | | | **PSI KO** | **Rorβ KO** |
| # | **F** | | 3/ 94 | 3/114 | 3/107 |
| # | **G** | | 3/ 94 | 3/114 | 3/107 |
| | **H** | | 3/1558 | 3/ 2453 | 3/ 2088 |

| Figure S4 | panel | sub-panel | Control (N/n) | Condition (N/n) | Condition (N/n) |
|---|---|---|---|---|---|
| | | | | **PV-silencing** | |
| | **A** | brush | 4/ 99 | 4/127 | |
| # | **B** | | 4/ 99 | 4/108 | |
| | | | 4/ 99 | 4/108 | |
| # | **C** | | 4/ 99 | 4/108 | |
| | **D** | | 4/1410 | 4/1458 | |
| | | | **Sham shuffled (n)** | **SNI shuffled (n)** | **PV-silencing (N=4)** |
| # | **E** | indentation | 125 | 125 | 108 |
| | | brush | 125 | 125 | 127 |
| # | **F** | superficial | 30 | 30 | 24 |
| | | deep | 100 | 100 | 84 |
| | **G** | noise | 1400 | 1400 | 1387 |
| | | indentation | 1800 | 1800 | 1797 |
| | | | **Control** | **PSI KO** | **Rorβ KO** |
| | **I** | noise | 3/1558 | 3/ 2453 | 3/ 2088 |
| | | indentation | 3/1558 | 3/ 2453 | 3/ 2088 |

| Figure | panel | test | statistic | p-value |
|---|---|---|---|---|
| 1 | D | Mann-Whitney U | 109679 | 0.006 |
| 2 | A | Mann-Whitney U | 114728 | <0.0001 |
| | B superficial | Mann-Whitney U | 4565 | 0.0016 |
| | C | Mann-Whitney U | 30879621 | <0.0001 |
| | D | Mann-Whitney U | 31474258 | <0.0001 |
| | E | Mann-Whitney U | 30076950 | <0.0001 |
| 3 | G | Mann-Whitney U | 76 | 0.0322 |
| | H | Unpaired t test | t=2.705, df=29 | 0.0113 |
| 4 | A | Mann-Whitney U | 2533 | <0.0001 |
| | B superficial | Mann-Whitney U | 135 | 0.0071 |
| | C | Mann-Whitney U | 765752 | <0.0001 |
| | D noise | Mann-Whitney U | 913091 | 0.0024 |
| | D signal | Mann-Whitney U | 789032 | <0.0001 |
| | I | Mann-Whitney U | 0.5 | 0.0043 |
| S2 | A | Mann-Whitney U | 33004 | <0.0001 |
| | B 3ms | Mann-Whitney U | 106197 | <0.0001 |
| | C | Mann-Whitney U | 15222303 | <0.0001 |
| | D latency 75mN | Mann-Whitney U | 57131 | <0.0001 |
| | D jitter 10mN | Mann-Whitney U | 11532 | 0.0217 |
| | D jitter 75mN | Mann-Whitney U | 68612 | 0.0226 |
| | E indentation | Mann-Whitney U | 3256 | <0.0001 |
| | E brush | Mann-Whitney U | 6737 | <0.0001 |
| | F | Mann-Whitney U | 2103818 | <0.0001 |
| | G noise | Mann-Whitney U | 2447068 | <0.0001 |
| | G indentation | Mann-Whitney U | 2503596 | <0.0001 |
| | G brush | Mann-Whitney U | 3687353 | <0.0001 |
| S4 | A 3ms | Mann-Whitney U | 4222 | 0.0009 |
| | B | Mann-Whitney U | 3887 | <0.0001 |
| | D | Mann-Whitney U | 733833 | <0.0001 |

*all tests are two-tailed
